# Epidermal ZBP1 stabilizes mitochondrial Z-DNA to drive UV-induced IFN signaling in autoimmune photosensitivity

**DOI:** 10.1101/2024.01.23.576771

**Authors:** Benjamin Klein, Mack B. Reynolds, Bin Xu, Mehrnaz Gharaee-Kermani, Yiqing Gao, Celine C. Berthier, Svenja Henning, Shannon N. Loftus, Kelsey E. McNeely, Amanda M. Victory, Craig Dobry, Grace A. Hile, Feiyang Ma, Jessica L. Turnier, Johann E. Gudjonsson, Mary X. O’Riordan, J. Michelle Kahlenberg

## Abstract

Photosensitivity is observed in numerous autoimmune diseases and drives poor quality of life and disease flares. Elevated epidermal type I interferon (IFN) production primes for photosensitivity and enhanced inflammation, but the substrates that sustain and amplify this cycle remain undefined. Here, we show that IFN-induced Z-DNA binding protein 1 (ZBP1) stabilizes ultraviolet (UV)B-induced cytosolic Z-DNA derived from oxidized mitochondrial DNA. ZBP1 is significantly upregulated in the epidermis of adult and pediatric patients with autoimmune photosensitivity. Strikingly, lupus keratinocytes accumulate extensive cytosolic Z-DNA after UVB, and transfection of keratinocytes with Z-DNA results in stronger IFN production through cGAS-STING activation compared to B-DNA. ZBP1 knockdown abrogates UV-induced IFN responses, whereas overexpression results in a lupus-like phenotype with spontaneous Z-DNA accumulation and IFN production. Our results highlight Z-DNA and ZBP1 as critical mediators for UVB-induced inflammation and uncover how type I IFNs prime for cutaneous inflammation in photosensitivity.

**One Sentence Summary:** ZBP1 and mitochondrial Z-DNA drive autoimmune photosensitivity via cGAS-STING activation.

## INTRODUCTION

Autoimmune photosensitivity is seen in type I Interferon (IFN)-driven skin diseases such as systemic (SLE) and cutaneous lupus erythematosus (CLE) as well as dermatomyositis (DM) (*1–4*). Up to 81% of SLE and CLE patients are affected by photosensitivity, defined by severe skin reactions to brief ultraviolet light (UV). Photosensitivity leads to poor quality of life for patients(*1, 5*), and in SLE, UV light can also trigger systemic inflammation, including nephritis(*1, 6–8*). Therapies to prevent photosensitivity are limited to sunscreen use, for which compliance is low(*9*).

The skin is the most common affected organ of lupus patients(*10*). Nonlesional skin of patients at risk for lupus has been shown to exhibit robust IFN secretion even with minimal IFN signature in the blood(*10–12*). This cutaneous type I IFN production leads to increased epidermal cell death(*3*), inflammatory activation of myeloid cells(*12*), and is widely accepted as a driver of CLE and DM lesions(*3, 11, 13, 14*). Within the skin, basal keratinocytes (KCs) are the main target of UVB irradiation and are major contributors to the IFN signature observed in these diseases through chronic secretion of IFN kappa (IFNκ)(*3, 15, 16*). Like type I IFNs, type III IFNs (composed of IFNλ 1-4) are upregulated in CLE skin and may also impact inflammatory responses in photosensitivity(*17, 18*). It is assumed that the IFN signature in nonlesional skin of lupus patients enhances immune activation and thus contributes to the transition to lesional skin or even systemic disease after environmental triggers, such as UV light(*11*). However, the precise mechanism by which a cutaneous type I IFN-rich environment accelerates inflammation after UV light has not been identified.

The effects of UV light in KCs include mitochondrial DNA damage, mitochondrial fragmentation, and release of mtDNA into the cytoplasm(*19–21*). UV light is mainly absorbed by complex I of the mitochondrial electron transport chain and leads to reactive oxygen species (ROS) accumulation and disruption of respiratory chain complexes(*22*). MtDNA is especially susceptible to UV-induced damage as it is not protected by histones, is located in the inner mitochondrial matrix in close proximity to produced ROS, and lacks the same mechanisms that repair nuclear DNA(*19*). UV light causes type I IFN responses in the skin which is accelerated in autoimmune photosensitivity(*1, 2, 23*). It has been proposed that this IFN stems from cytoplasmic nucleic acids that activate innate immune sensors. Despite growing evidence that this happens in a cyclic GMP-AMP synthase (cGAS)-stimulator of interferon genes (STING)-dependent fashion(*24, 25*), the substrate of cGAS activation has not been identified after UV exposure.

MtDNA is a major activator of type I IFN responses in multiple autoimmune diseases including SLE(*26–30*). Specifically, oxidized mtDNA is highly interferogenic in lupus neutrophils(*28, 29*). Moreover, mtDNA derived from mitochondria-containing red blood cells activates IFN responses in lupus monocytes contributing to the IFN signature observed in SLE blood(*31*).

Mitochondrial dysfunction can lead to liberation of mtDNA into the cytoplasm(*27, 32, 33*). After release of mtDNA, multiple pattern recognition receptors (PRRs) including Toll-like receptor 9 (TLR9), cGAS, and Z-DNA binding protein 1 (ZBP1) can be activated(*26, 32–34*). ZBP1 itself represents an IFN-induced gene sensing nucleic acids in Z-conformation with its Zα domain to provide antiviral defense(*35–39*). In contrast to B-DNA, left-handed Z-DNA is more prone to occur in GC rich sequences and is characterized by a zig-zag shaped backbone. This DNA-conformation is induced by high salt conditions, torsional stress, oxidized bases such as 8-Oxo-2’-deoxyguanosine (8-oxodG) or other base modifications and further stabilized by Z-DNA binding proteins(*35, 40, 41*). MtDNA can undergo Z-DNA formation upon mitochondrial genome instability and negative supercoiling(*34*). Importantly, mitochondrial-derived Z-DNA is detected by ZBP1 and cGAS to sustain type I IFN-responses in cardiomyocytes to promote doxorubicin-induced cardiotoxicity(*34*).

Given mitochondrial damage after UV light and the type I IFN signature in autoimmune photosensitive diseases, we hypothesized that UV exposure could lead to oxidized mitochondria-derived cytosolic Z-DNA accumulation that is stabilized by type I IFN-induced upregulation of ZBP1. This combination of Z-DNA accumulation and increased ZBP1 would then lead to robust type I IFN responses in KCs. Here, we demonstrate that ZBP1 is upregulated in nonlesional and lesional skin of SLE and DM patients. We provide evidence of enhanced ZBP1-stabilized Z-DNA accumulation in an IFN-rich environment, which further sustains IFN signaling in KCs after UV exposure via cGAS-STING activation. Knockdown of ZBP1 in KCs attenuates type I IFN and IFN-stimulated gene (ISG) expression after UV, and overexpression of ZBP1 in KCs results in enhanced cytosolic Z-DNA retention and type I IFN signaling. Together, our data identify a novel pathway that explains how an IFN-rich environment primes the skin for photosensitivity through ZBP-1-mediated Z-DNA sensing which drives activation of the cGAS-STING pathway. This has important implications for treatment and prevention of cutaneous and systemic flares of photosensitive autoimmune diseases.

## RESULTS

### Ultraviolet irradiation induces mtROS-dependent type I and III IFN induction in KCs accompanied by cytosolic Z-DNA accumulation derived from mitochondria

UV light was previously shown to cause mitochondrial damage and mitochondrial ROS (mtROS) accumulation (*22*). However, the downstream effects of these changes have not been well-characterized.

We hypothesized that UV-induced mtROS and release of mtDNA would lead to IFN production in KCs. To test this, we irradiated N/TERT keratinocytes with UVB light and preincubated them with or without mitoTEMPO, a mitochondrially targeted antioxidant (Fig. 1,A). We observed, via mitoSOX Red staining, increased mtROS formation in KCs 30min after UVB exposure that was inhibited by mitoTEMPO (Fig. 1,B and C). To test whether mtROS promote type I and III IFN signaling after UV light, we assessed the effect of mitoTEMPO on type I and III IFN gene expression. Rotenone, a complex I inhibitor and inducer of mtROS(*42*), was used as a positive control. Both UVB and Rotenone induced a significant increase in *IFNB1* and *IFNL3* expression 6h after UVB which was rescued by mitoTEMPO (Fig. 1,D). MitoTEMPO also dampened expression of later expressed genes, *IFNK, OASL* and *MX1* 24 hours after UV exposure (Fig. 1,E). These results indicate that mtROS are promoting type I and III IFN responses after UV in KCs. To test whether mtDNA is required for UV-induced type I IFN induction, we then depleted mtDNA selectively with the nucleoside 2’3’-dideoxycytosine (ddC) and observed a reduction of type I IFN in a dose dependent manner (Supplemental Fig. 1,A and B). This suggests that mtROS effects on mtDNA are important for UV-driven IFN production.

**Figure 1.**
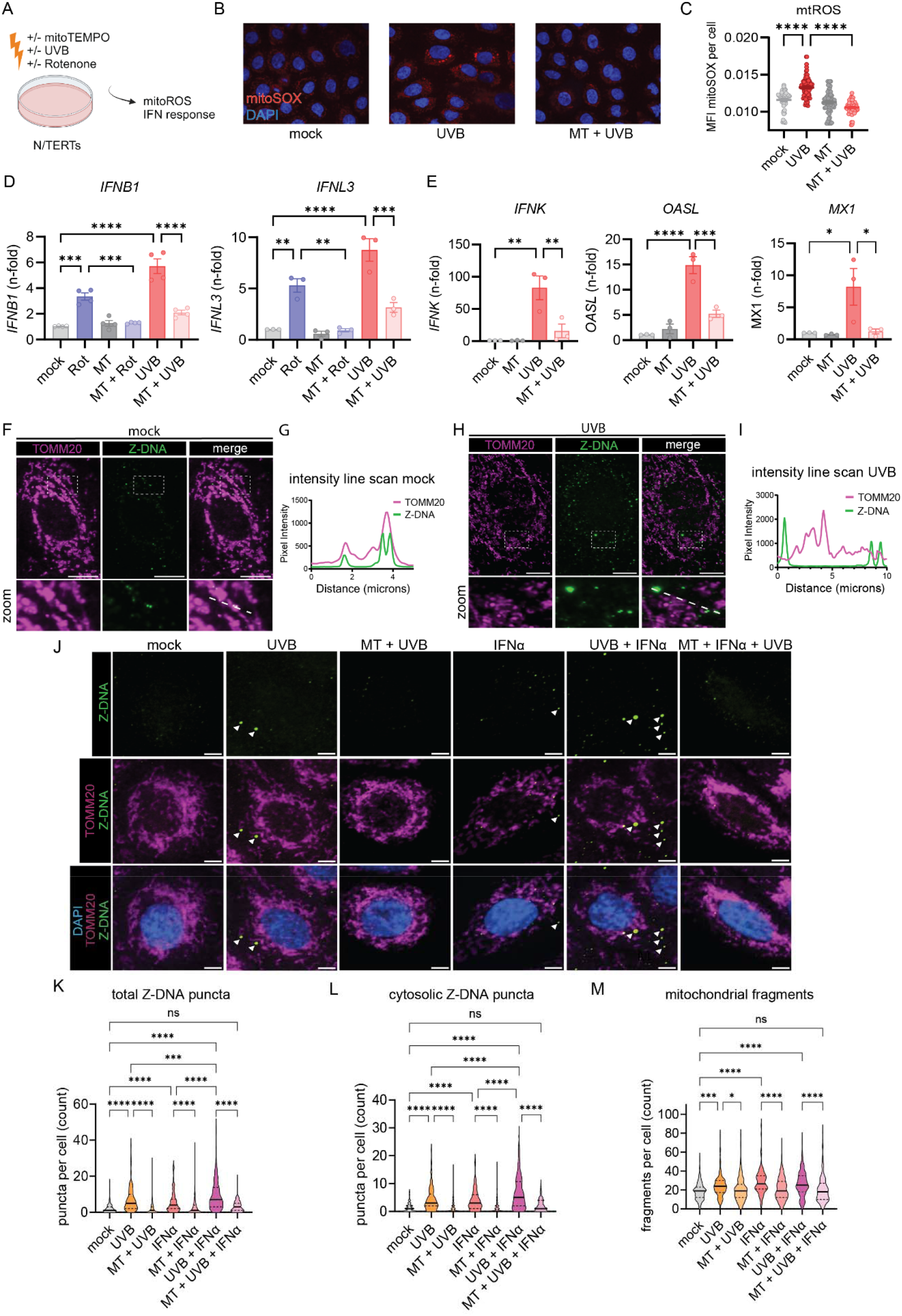
UVB light causes mtROS dependent IFN responses accompanied by cytosolic Z-DNA release derived from mitochondria. **A.** Experimental approach. **B.** Representative images from N/TERTs treated +/-mitoTEMPO (MT) +/-UVB irradiation stained with MitoSOXred and Hoechst33342. **C.** Quantification of MitoSOX intensity per cell using CellProfiler software. **D.** N/TERTs were treated with either rotenone, MT or UVB for 6h. Gene expression of *IFNB1* (n=44) and *IFNL3* (n=3) was determined by qPCR. **E.** Gene expression of *IFNK*, *MX1* and *OASL* (n=3) +/-mitoTEMPO +/-UVB 24h after UVB exposure. **F.** Representative confocal microscopy from N/TERTs of TOMM20, Z-DNA, and DAPI without stimulation. **G.** Line scan analysis of the line in F. **H.** Representative confocal microscopy from N/TERTs of TOMM20 and Z-DNA 3h after UVB exposure. **I.** Line scan analysis of the dotted white line in H. **J.** Representative confocal images from N/TERTs of TOMM20, Z-DNA, DAPI +/-mitoTEMPO, pretreatment with IFNα or 3h after UVB. Scale bar 5μm. **K-L.** Quantification of Z-DNA puncta using CellProfiler software. **M.** Mitochondrial fragments (objects <1µm^2^ with circularity > 0.6) using CellProfiler. Mean + SEM or violin plots with mean + quartiles of n≥3 independent experiments. P-values were calculated using ordinary one-way ANOVA followed by Sidak’s multiple comparison test. *P < 0.05; **P < 0.01; ***P < 0.001; ****P < 0.0001.

Oxidation of DNA and mitochondrial DNA instability can lead to formation and stabilization of left-handed Z-DNA(*34, 40*). Therefore, we assessed Z-DNA localization and accumulation after UV exposure using quantitative immunofluorescence microscopy of Z-DNA, the mitochondrial outer membrane protein TOMM20 and DAPI. Utilizing a Z22 antibody which was confirmed to stain Z-DNA in previous reports(*34*), we screened for enhanced Z-DNA staining and accumulation after UVB exposure. Using super-resolved structured illumination microscopy, we observed low baseline staining and mitochondrial localization of Z-DNA without stimulation (Fig. 1,F and G). Strikingly, after UVB we observed an increase of Z-DNA together with translocation from the mitochondrial compartment into the cytoplasm with formation of prominent Z-DNA puncta (Fig. 1,H and I). Further using a spinning disk confocal microscope to image larger cell numbers, we observed prominent Z-DNA puncta whereas small puncta were comparatively weaker (Fig. 1,J). Automated image analysis revealed that UVB significantly increased total and cytosolic Z-DNA 3h after irradiation which was rescued by preincubation with mitoTEMPO (Fig. 1,K and L). Moreover, we observed significantly more mitochondrial fragments after UVB exposure which was also rescued by mitoTEMPO (Fig. 1,M). Z-DNA was localized within the mitochondrial network when cells were preincubated with mitoTEMPO (Fig. 1J), suggesting a stabilization of the mitochondrial network and inhibition of Z-DNA accumulation through scavenging of mtROS. We confirmed mitochondrial origin of Z-DNA by depletion of mtDNA using ddC, which reduced Z-DNA intensity and puncta significantly but did not influence UVB-induced mitochondrial fragmentation (Supplemental Fig. 1D-F).

Type I IFNs have been implicated in photosensitive responses and are responsible for an autocrine loop of inflammation upon UV exposure(*3, 13*). To explore whether type I IFNs impacted the UVB effects on mitochondrial stress and cytosolic Z-DNA accumulation, we first treated N/TERTs for 16h with IFNα before irradiation to mimic a chronic type I IFN environment, such as seen in SLE(*3, 10–12*). Via microscopy, we observed an increase in cytosolic Z-DNA puncta and mitochondrial fragmentation in N/TERTs after IFNα priming alone (Fig 1. K, L). After UVB exposure and IFNα priming, we observed a striking accumulation of cytosolic Z-DNA puncta associated with mitochondrial fragmentation (Fig. 1,L and M). MitoTEMPO fully rescued this phenotype by maintaining the mitochondrial network (Fig. 1,M). Collectively, these results strongly indicate cytosolic Z-DNA accumulation derived from mitochondria upon UV-exposure is dramatically increased in a type I IFN rich environment.

### UVB-induced oxidative DNA damage promotes cytosolic Z-DNA accumulation

Given our observations of UVB-induced mtROS and the reduction of Z-DNA by mitoTEMPO, we suspected that UVB-driven oxidative mtDNA lesions would contribute to Z-DNA accumulation. Previous published evidence showed that oxidative damage of Guanosine (8-Oxo-2’-deoxyguanosine, 8oxodG) promotes Z-DNA formation through energetic favorability, as 8oxodG hinders steric formation of B-DNA and favors Z-DNA formation(*40*). Staining for 8oxodG revealed cytosolic increases upon UVB exposure whereas these lesions were partially prevented by mitoTEMPO (Supplemental Fig. 1, C and D). Importantly, while 8-oxodG staining was diffuse, co-staining with Z-DNA and TOMM20 revealed proximity of large Z-DNA puncta with spots of enhanced 8oxodG damage outside of mitochondria after UVB exposure (Fig. 2,A and B). After IFNα priming and UVB exposure, we observed that accumulation of large Z-DNA puncta occurred in spots with increased 8oxodG staining (Fig. 2,A-C). There was a significant positive correlation of cytosolic Z-DNA puncta and 8oxodG intensity per cell, suggesting 8oxodG damage promotes Z-DNA conformation (Fig. 2,D). In addition, mitochondrial fragmentation after IFNα priming and UVB exposure exhibited a similar positive correlation with cytosolic Z-DNA accumulation (Supplemental Fig. 3,A-E). These results suggest cytosolic Z-DNA accumulation is associated with mitochondrial fragmentation and oxidative DNA damage.

**Figure 2.**
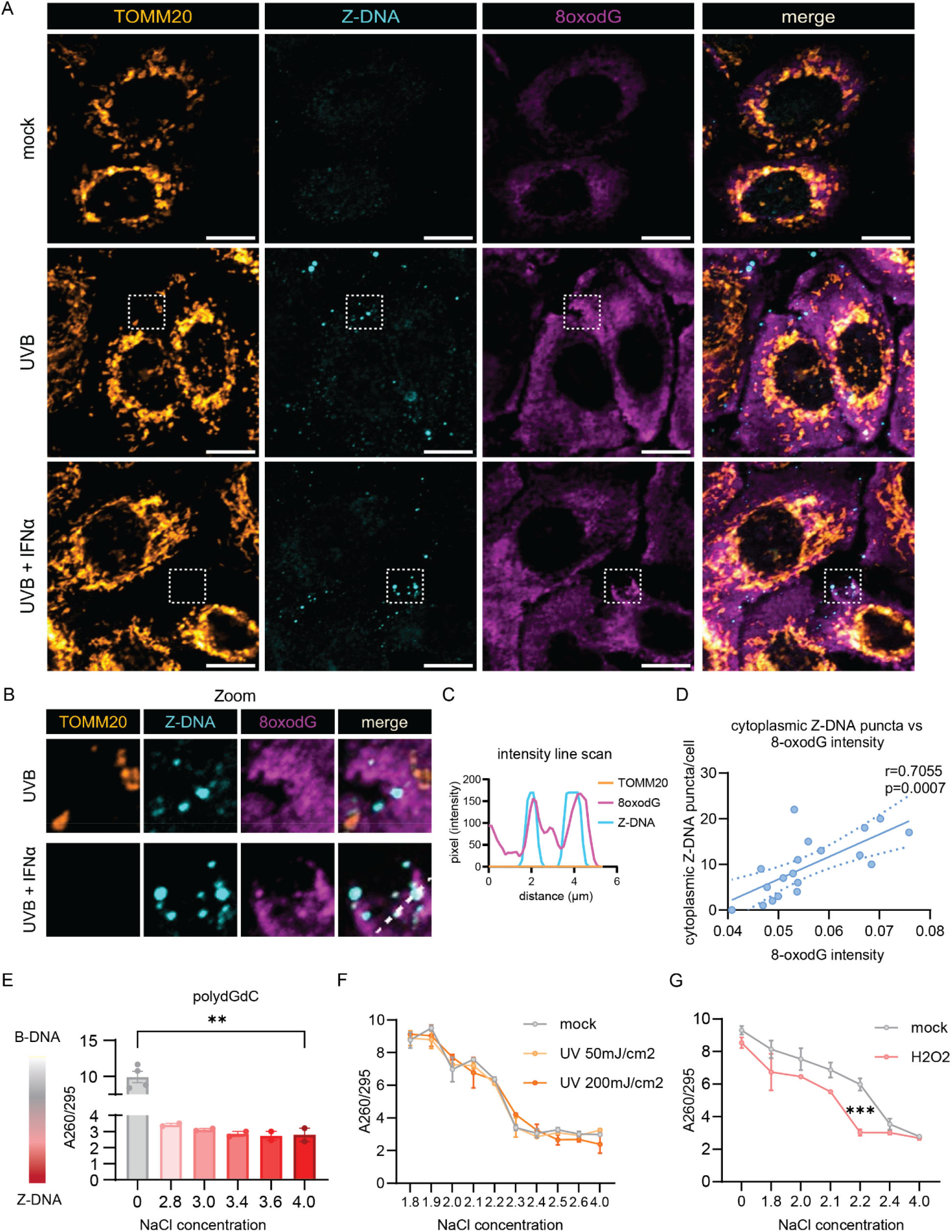
UVB promotes Z-DNA formation via oxidative DNA damage. **A.** Representative confocal images of the mitochondrial outer membrane (TOMM20), Z-DNA and 8oxodG in N/TERTs at baseline, 3h after UVB exposure with or without IFNα preincubation for 16h. Scale bar 10μm. **B.** Magnified region from (A) highlighting proximity and colocalization of Z-DNA with intense 8oxodG staining in areas outside of mitochondria. **C.** Representative line scan from Z-DNA puncta in (B) highlighting the absence of TOMM20 in spots of Z-DNA accumulation. **D.** Correlation of the Z-DNA puncta per cell with matched average 8oxodG per cell. **E.** Change of A260/295 as a measure of B-DNA (high ratio∼10) vs Z-DNA (lower ratio of ∼3) formation is graphed comparing low salt vs. high salt conditions after 2h at 37°C. **F.** Naked polydGdC was irradiated with indicated UVB doses and incubated in indicated [NaCl] as in E. to induce Z-DNA. No shift to a lower ratio in lower [NaCl] was detected after UVB light exposure. **G.** To test the effect of oxidation on propensity for Z-DNA formation, polydGdC was treated with H2O2 (1mM) for 2h at 37°C and subjected to varying salt concentrations as in E. Buffers with indicated NaCl and H2O2 without DNA served as blanks for the assay.

To test whether Z-DNA formation required oxidative lesions, we utilized an assay that examines Z-DNA formation in polydGdC oligos induced by high [NaCl](*43, 44*). Z-DNA conformation under high salt was confirmed using the A260/295 ratio which is significantly lower in Z-DNA compared to the B conformation (Fig. 2,E)(*43, 45*). To test whether UVB exposure would directly lead to Z-DNA accumulation, we irradiated polydGdC with low (50mJ/cm^2^) and high (200mJ/cm^2^) doses of UVB light. We did not observe direct induction of Z-DNA conformation upon UVB exposure (Fig. 2,F), suggesting no direct oxidative DNA damage at the doses used in our *in vitro* system. However, induction of oxidation by H_2_O_2_ permitted Z-DNA formation at significantly lower concentrations of NaCl (Fig. 2,G). These results suggest that UVB light promotes Z-DNA conformation indirectly by accumulation of oxidative DNA damage. We then examined whether IFNα would increase cytoplasmic Z-DNA by augmenting mitochondrial or cellular ROS. Surprisingly, IFNα did not enhance ROS generation, suggesting that IFNα-induced increase of cytoplasmic Z-DNA accumulation is independent of ROS generation through IFNα (Supplemental Fig. 4,A and B).

### ZBP1 is overexpressed in photosensitive autoimmune skin diseases and correlates with type I IFN scores

Autoimmune photosensitive diseases share a cutaneous IFN signature in lesional and non-lesional skin that can be further induced after UV exposure to promote chronic cutaneous inflammation(*1, 2, 10–13, 25*). As UVB-mediated cytosolic mtDNA in Z form is promoted by type I IFNs (Fig. 1, J-L), we hypothesized that the chronic type I IFN high environment in autoimmune photosensitive diseases may contribute to this sustained inflammation. Intriguingly, Z-DNA conformation can be stabilized by the IFN-regulated gene Z-DNA binding protein 1 (*ZBP1*)(*34–36*), so we chose to further examine the role of ZBP1 in autoimmune photosensitive diseases (Fig. 3,A). Strikingly, *ZBP1* expression was significantly upregulated in a large cohort of CLE lesional skin biopsies (n=90) compared to healthy controls (CTL) (n=13) (Fig. 3,B). This upregulation was independent of CLE subtype and presence or absence of systemic disease (Fig. 3,B). Furthermore, *ZBP1* expression correlated significantly with IFN scores in CLE, suggesting its upregulation to be a phenomenon of cutaneous IFN expression rather than specific systemic clinical features (Fig. 3,C). We confirmed these results in microarray datasets from patients with childhood SLE (cSLE) (n=7), where we observed significant upregulation of *ZBP1* gene expression compared to CTL (n=8) and significant correlation with the cutaneous IFN score (Fig. 3,D and E). To assess nonlesional *ZBP1* expression in the skin and identify which cells are expressing *ZBP1*, we utilized our single-cell-sequencing dataset of lupus patients (n=14) compared to CTL (n=14)(*12*). We confirmed robust upregulation of *ZBP1* in nonlesional and lesional keratinocytes compared to CTL (Fig. 3,F). There was also a similar, but weaker, trend in cutaneous myeloid cells and endothelial cells; *ZBP1* expression was mostly absent in fibroblasts, melanocytes, and mast cells (Fig. 3,F).

**Figure 3.**
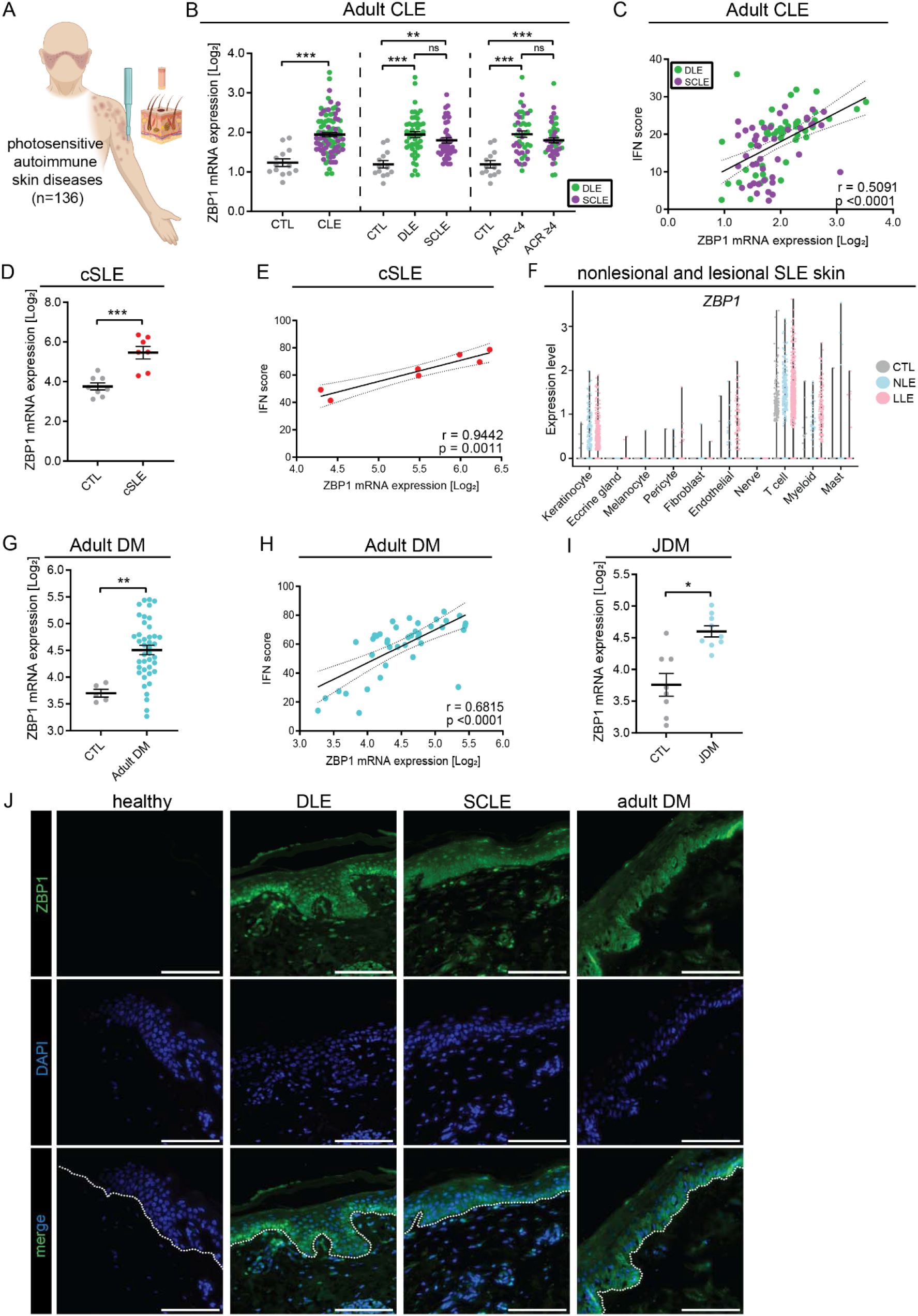
ZBP1 is overexpressed in the epidermis of autoimmune photosensitive diseases. **A.** Graphical representation of data acquisition from lesional skin microarrays. **B.** *ZBP1* expression in lesional cutaneous lupus (CLE) (n=90) compared to healthy control (CTL) (n=13) (left), by lesion subtype (discoid lupus erythematosus (DLE) or subacute cutaneous lupus erythematosus (SCLE), middle) and based on the presence or absence of systemic lupus via >4 1997 ACR criteria (right). **C.** Correlation of *ZBP1* expression in CLE with IFN score, linear regression. **D.** *ZBP1* expression in childhood onset systemic lupus erythematosus (cSLE, n=7) compared to CTL (n=8). **E.** Correlation of ZBP1 expression with IFN score in cSLE. **F.** Violin plots showing *ZBP1* expression from scRNA sequencing across cutaneous cell types from nonlesional lupus skin (NLE, n=14), lesional lupus skin (LLE, n=14) compared to CTL (n=14). **G.** Expression of *ZBP1* in adult dermatomyositis (DM) (n=41) and **H.** Correlation with IFN score by linear regression. **I.** *ZBP1* expression in juvenile dermatomyositis (jDM, n=9) compared to CTL (n=8). **J.** Representative images of tissue imunofluoresence of ZBP1 in CTL (n= 7), DLE (n=8), SCLE (n=5) and DM (n=6). Dotted white line indicates the dermo-epidermal junction. Scale bar =100μm. Mean + SEM. * = q <D0.05; ** = q <D0.01; *** = q <D0.0001, by Student’s unpaired t-test.

Further analysis of adult DM (n=30) and JDM (n=9) compared to CTL (n=8 and n=5, respectively) revealed significant cutaneous *ZBP1* upregulation in DM and JDM with significant correlation of *ZBP1* expression with the IFN score in adult DM (Fig. 2,G-I). In JDM, we were underpowered to observe a significant correlation between IFN score and *ZBP1*, but there was a similar trend (Supplemental Fig. 5,A). The expression of *ZBP1* in lupus skin samples neither correlated with autoantibody status, disease activity nor with age (Supplemental Fig. 5, B and C), suggesting upregulation of *ZBP1* is independent of these patient characteristics.

Next, we examined the protein expression of ZBP1 by staining skin samples from nonlesional and lesional lupus and DM skin. As expected, we observed increased ZBP1 in nonlesional and lesional discoid lupus erythematosus (DLE), subacute cutaneous lupus eyrthematous (SCLE) and DM epidermis compared to CTL, where ZBP1 was basically undetectable (Fig. 3,J). Notably, ZBP1 was highest expressed in the basal layer of the epidermis. In sum, these results indicate epidermal upregulation of ZBP1 in nonlesional and lesional skin of autoimmune photosensitive diseases correlates with cutaneous IFN signatures and suggests that ZBP1 may be important for downstream immune responses in adult and pediatric CLE, SLE and DM skin.

### Lupus KCs exhibit strong baseline and UV-induced cytosolic Z-DNA compared to CTL

Given enhanced IFN signaling and significant upregulation of ZBP1 in lupus KCs, we quantified cytosolic Z-DNA puncta in CTL and lupus KCs at baseline and after UVB exposure (Fig. 4,A). At baseline, CTL KCs exhibited Z-DNA staining primarily within the mitochondrial network (Fig. 4,A). After UVB exposure, we observed the expected cytosolic Z-DNA accumulation in CTL KCs (Fig. 4,A and B). Strikingly, lupus KCs exhibited enhanced total and cytosolic Z-DNA puncta at baseline that were significantly increased after UVB with formation of multiple large cytosolic puncta (Fig 4,A and B). Preincubation of CTL KCs with IFNα promoted enhanced cytosolic Z-DNA accumulation after UVB exposure, comparable with nonlesional lupus KCs after UVB alone, suggesting that the known chronic IFN loop in SLE KCs(*3, 13*) stabilizes Z-DNA, potentially through the ISG ZBP1. Surprisingly, Z-DNA accumulation in the cytosol after UVB exposure of lupus KCs was prevented by preincubation with mitoTEMPO, which led to stabilization of Z-DNA within the mitochondrial network (Fig. 4,C).

**Figure 4.**
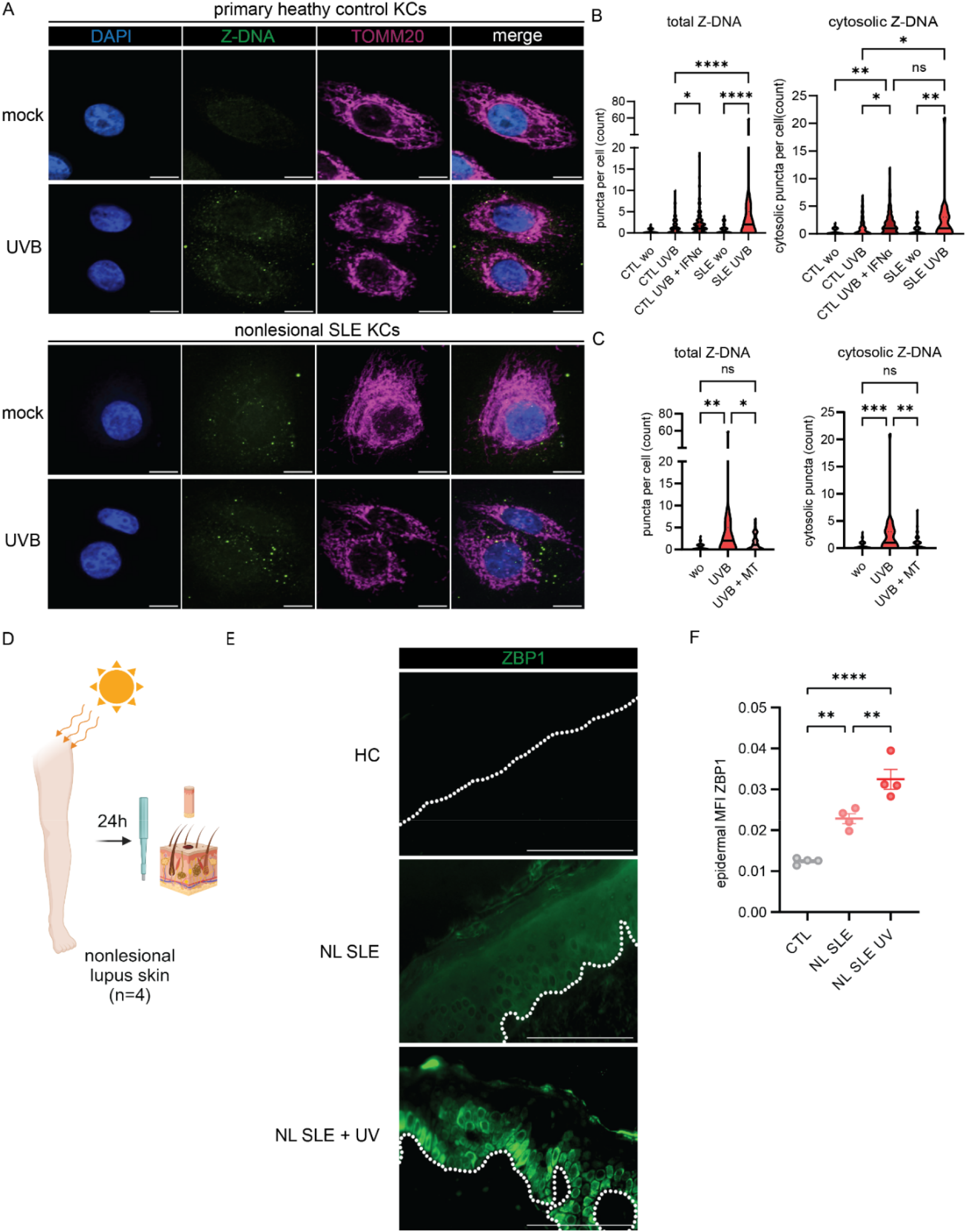
Nonlesional lupus keratinocytes exhibit cytosolic Z-DNA accumulation at baseline and after UVB exposure that is prevented by mitoTEMPO. **A.** Representative images of confocal microscopy staining for Z-DNA, TOMM20 and counterstaining with DAPI at baseline and after UVB exposure and preconditioning with IFNα (1000U/ml for 16h prior to UVB exposure) in primary healthy control KCs (n=4) and SLE KCs (n=3). **B-C.** Quantification of total and cytosolic Z-DNA puncta after UVB with or without preincubation with mitoTEMPO or IFNα using CellProfiler. **D.** Healthy controls (HC), SLE patients +/-UVB (n=4 each group) were biopsied 24h after UV exposure. **E.** Representative images of ZBP1 staining in HC, nonlesional SLE skin (NL SLE) and NL SLE after UV exposure. Dotted white line indicates the dermo-epidermal junction. Scale bar =100μm **F.** Quantification of mean fluorescence intensity (MFI) of epidermal ZBP1 using open source CelProfiler software. Ordinary one–way ANOVA followed by Sidak’s multiple comparison test. Mean and SEM. *P<0.05, **P<0.01, ***P<0,001, ****P<0.0001.

As UVB treatment of KCs upregulates type I and type III IFNs, we examined whether mitoTEMPO also decreased UV-induced IFN gene expression in lupus primary KCs. In line with our microscopy data, we observed that mitoTEMPO led to significant downregulation of *IFNB1, IFNL3* and ISG*s* after UVB exposure (Supplemental Fig. 6,A and B). Baseline and UVB-induced cellular ROS were not increased in SLE KCs compared to primary HC KCs (Supplemental Fig. 6,C). To further assess the role of ZBP1 *in vivo*, we assessed ZBP1 protein expression in lupus nonlesional skin samples collected by biopsy 24 hours after irradiation with a minimal erythema dose of UVB (Fig. 4,D). Strikingly, in addition to upregulation of ZBP1 in nonlesional SLE skin compared to CTL, we observed enhanced expression and cytosolic localization of ZBP1 in the basal epidermis of nonlesional SLE skin after UV exposure (Fig. 4,E and F), suggesting a role for ZBP1 in UV-mediated responses in SLE *in vivo*. Together, these results show activation of the Z-DNA/ZBP1 pathway in SLE compared to HC skin after UV exposure.

### Z-DNA has stronger immunostimulatory properties than B-DNA to sustain STING-dependent IFN responses in human KCs

ZBP1 was recently described to bind cGAS within the cytoplasm resulting in bridging of Z-DNA to cGAS and subsequent STING activation and sustained type 1 IFN responses(*34*). Hence, we next assessed the effects of ZBP1 and DNA in the Z conformation on IFN activation in KCs. First, we confirmed upregulation of *ZBP1* after IFNα treatment in N/TERTs and primary KCs (Fig. 5,A and B). We then assessed co-localization of Z-DNA and ZBP1 after UVB with and without IFNα priming. As shown in Fig. 5C, UV treatment induced co-localization of Z-DNA puncta and ZBP1 in the cytoplasm, and this was expectedly enhanced when cells were first primed with IFNα. These data confirmed previous study results that ZBP1 and Z-DNA form a cytosolic complex and that type I IFNs enhance this through upregulation of ZBP1 and increased Z-DNA formation(*34*).

**Figure 5.**
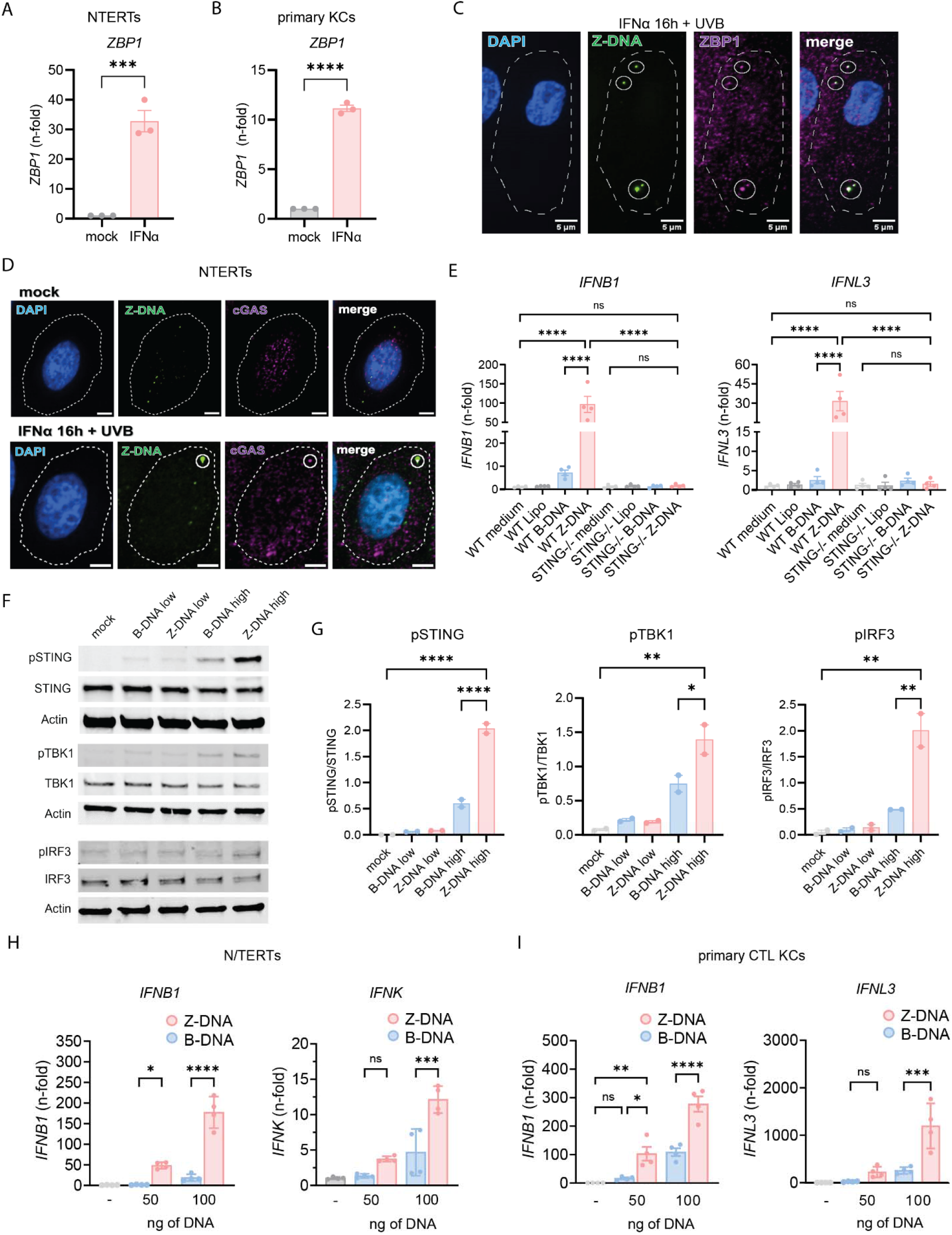
Z-DNA binds to ZBP1, activates the cGAS-STING pathway and has stronger immunostimulatory properties than B-DNA. **A-B.** Gene expression of *ZBP1* after IFNα stimulation in N/TERTs **(A.)** and primary healthy control KCs **(B.)** compared to β-Actin. **C.** Representative image of confocal microscopy from N/TERTs preincubated with IFNα (1000U/ml) and then irradiated with UVB exposure stained for Z-DNA, ZBP1, and DAPI 3h after UVB exposure. Scale bar 5μm. **D.** Representative images of IFNα-treated N/TERTs stained for Z-DNA, cGAS and DAPI 3h after UVB exposure. Cellular outline was drawn based on CellTrackerRed counterstain. **E.** Gene expression at 6h of indicated genes from N/TERTs and STING KO N/TERTs treated with Lipofectamine2000 alone or transfected with Z-DNA or B-DNA (500ng/ml). **F.** Representative Western Blot (n=2) of indicated proteins from N/TERTs transfected with 50ng (low) or 500ng (high) of Z-DNA (polydGdC) or B-DNA using Lipofectamine 2000. Lysates harvested 4h after transfection. **G.** Quantification of the abundance of pSTING, pIRF3, and pTBK1 relative to unphosphorylated proteins in transfected KCs. **H.** N/TERTs were transfected as in E and gene expression was measured 6h after DNA transfection (n=3). **I.** Primary control KCs (n=3) were transfected and analyzed as in H. Unpaired t-test and Ordinary one–way ANOVA followed by Sidak’s multiple comparison test. Mean and SEM. *P<0.05, **P<0.01, ***P<0,001, ****P<0.0001.

cGAS-dependent type I IFN responses have been described after UVB(*24*). Small Z-DNA puncta, which were primarily localized within the mitochondria (Fig. 1,F), did not colocalize with cGAS (Fig. 5,D). After UVB, we observed both cytoplasmic localization of cGAS as well as colocalization with large Z-DNA puncta (Fig. 5,D, Supplemental Fig. 7,A and B). This suggests that cGAS joins the complex of ZBP1 and Z-DNA within the cytosol after UVB exposure. To test whether Z-DNA signaling is cGAS-STING dependent, we transfected WT and STING^-/-^ N/TERTs with polydGdC, which is known to form Z-DNA conformation upon binding to positively charged Lipofectamine(*46*) and compared it to transfection of an equal amount of B-DNA. Both *IFNB1* and *IFNL3* expression were completely abrogated in STING^-/-^ KCs compared to mock knockout N/TERTs (Fig. 5,F). Intriguingly, we observed stronger activation of *IFNB1* and *IFNL3* expression 6h after Z-DNA transfection compared to B-DNA in WT KCs. We confirmed these results by Western Blot of pSTING, pTBK1 and pIRF3, revealing robust activation of this pathway by Z-DNA transfection (Fig. 5,G and H). We then assessed type I and type III IFN production in response to B- or Z-DNA and identified a striking difference in the upregulation of IFN genes and ISGs in Z-DNA transfected KCs (Fig. 5,J, Supplemental Fig. 8,A and B). This was confirmed in primary CTL KCs (Fig. 5,K). Interestingly, *ZBP1* expression was also induced to a significantly greater extent by Z-DNA compared to B-DNA transfection (Supplemental Fig. 8,A and B).

Together, these results confirm interaction of Z-DNA with ZBP1 in human primary keratinocytes and identify Z-DNA as more immunostimulatory compared to B-DNA in a cGAS-STING dependent fashion.

### ZBP1 is required for IFN signaling after UVB exposure and its upregulation recapitulates an autoimmune photosensitive phenotype in KCs

Given upregulation of ZBP1 by IFNα and stronger Z-DNA accumulation after UVB in IFNα-primed KCs, we next wanted to explore whether knockdown of ZBP1 reduced UVB-induced type I IFN responses, especially in an IFN-rich environment. We thus generated a knockdown of *ZBP1* using shRNA in N/TERTs (Fig. 6,A, Supplemental Fig. 9,A). After UV exposure, KCs deficient in ZBP1 showed less type I and III IFN activation compared to controls (Fig. 6,B). Priming with IFNα prior to UVB exposure led to significant upregulation of *IFNB1* and *IFNL3* gene expression in shcontrol KCs whereas this enhancement was absent in shZBP1 KCs (Fig. 6,B). The robust upregulation of the ISG *CCL5* expression following priming with IFNα prior to UV was also dramatically decreased in the absence of ZBP1 (Fig. 6,C). These results suggest that ZBP1 is required for UV-driven IFN responses and is particularly crucial for the effects of type I IFN priming on cGAS/STING activation and IFN production.

**Figure 6.**
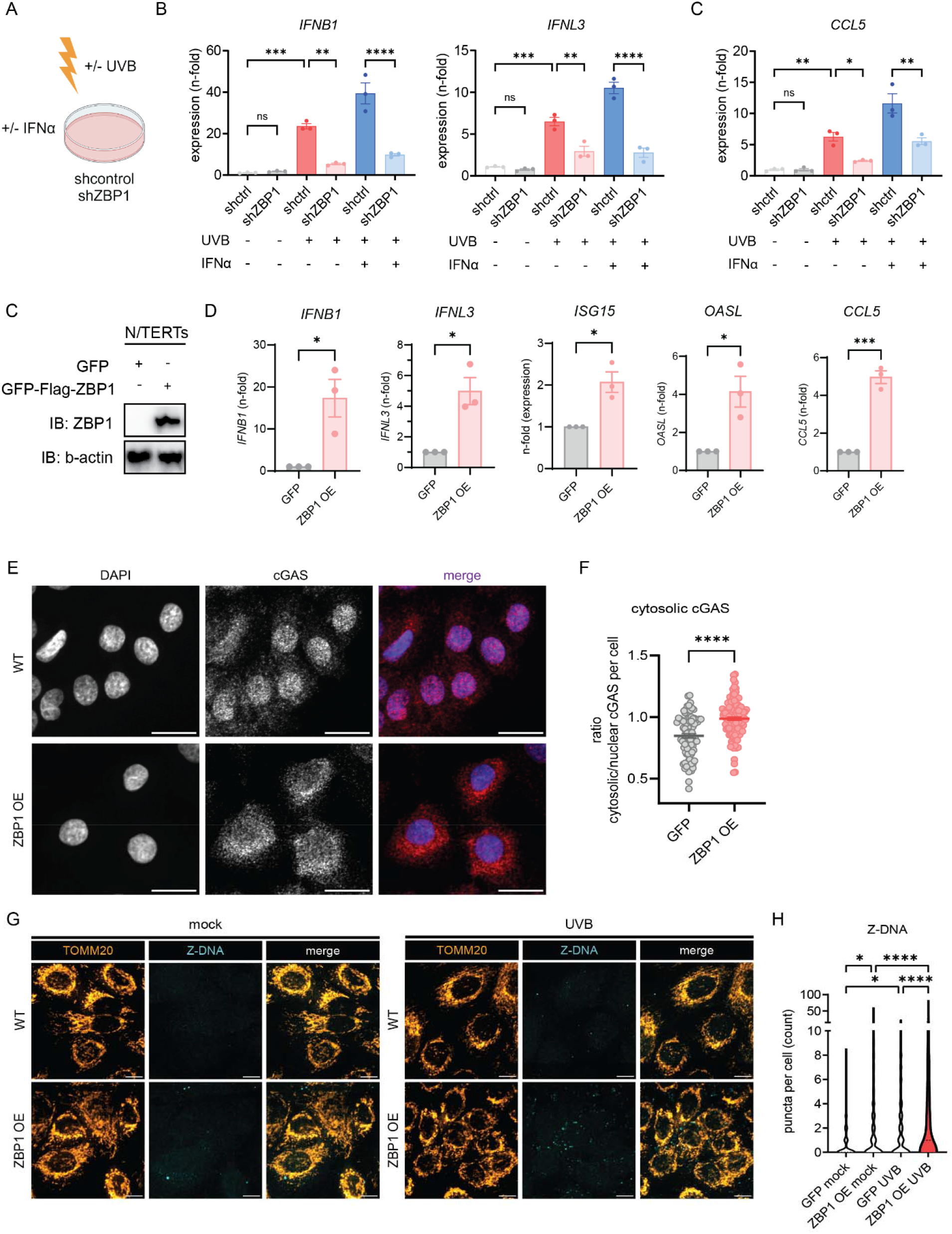
ZBP1 regulates UVB-induced type I and III IFN responses in human keratinocytes. **A-B.** Knockdown of ZBP1 in N/TERTs was performed using a lentivirus expressing either shRNA targeting Human ZBP1 or shcontrol. Baseline, UV, and UV+IFNα induced gene expression were assessed for **(A.)** Type I and III IFN (6h after UVB) and **(B.)** CCL5, a known ISG (24h after UVB); n=3 **C.** Generation of GFP/3XFLAG-tagged N/TERTs with overexpression of ZBP1 compared to GFP-tag only. **D.** IFN genes and ISGs were assessed by RT-qPCR at baseline in ZBP1 overexpressing cells (ZBP1 OE) or GFP-expressing control N/TERTs (GFP). **E.** Representative greyscale and merged images of confocal microscopy from N/TERTs with GFP or GFP-ZBP1 N/TERTs for cGAS (red) and DAPI (blue). Scale bar 20μm. **F.** Quantification of cytosolic cGAS using ratio of cGAS MFI in the cytosol versus nuclear cGAS MFI using CellProfiler. **G.** Representative confocal images of GFP-tagged N/TERTs or ZBP1 overexpressing N/TERTs of TOMM20 and Z-DNA at baseline and 3h after UVB exposure. Scale bar 10μm. Ordinary one –way ANOVA followed by Sidak’s multiple comparison test. Mean and SEM. *P<0.05, **P<0.01, ***P<0,001, ****P<0.0001.

ZBP1 is increased in the epidermis of adult and childhood SLE as well as in adult and juvenile DM. To mimic the phenotype observed in these autoimmune photosensitive diseases, we generated a GFP-ZBP1-FLAG tagged KC cell line in N/TERTs and compared it to a N/TERT line expressing GFP alone (Fig. 6,D, Supplemental Fig. 9, B and C). ZBP1 was primarily expressed in the cytoplasm in the overexpressing cell line (Supplemental Fig. 9,D). Upregulation of ZBP1 was confirmed by Western Blot (Fig. 6,D). We next tested for IFN gene expression in GFP-ZBP1-FLAG KCs. Strikingly, we observed significant upregulation of both type I and III IFNs as well as ISGs in GFP-ZBP1-FLAG KCs compared to GFP KCs without additional stimulation (Fig. 6,E). In addition, confocal microscopy revealed cytosolic localization of cGAS in ZBP1 overexpressing KCs, consistent with its activation (Fig. 6,G). We then wanted to know whether overexpression of ZBP1 was sufficient to increase cytosolic Z-DNA at baseline and after UV exposure. Indeed, cytosolic Z-DNA was increased at baseline in the ZBP1 overexpressing KCs compared to controls (Fig. 6,H and I). These results are in line with our previous observations that IFNα treatment alone can increase cytosolic Z-DNA (Fig. 1,J-L). After UVB exposure, we observed massive cytosolic Z-DNA accumulation in GFP-ZBP1-FLAG KCs compared to GFP control (Fig. 6,H and I). Importantly, mitochondrial fragmentation did not differ between GFP-ZBP1-FLAG KCs compared to GFP control (Supplemental Fig. 9, E). These data suggest that ZBP1 overexpression is sufficient to promote Z-DNA stabilization and its downstream IFN signaling but does not affect mitochondrial health per se.

Together, these results indicate that upregulation of ZBP1 is sufficient to stabilize Z-DNA and promote STING activation. Importantly, ZBP1 is required for pro-inflammatory effects of type I IFNs on UV-mediated STING activation. Upregulation of ZBP1 recapitulates the phenotype observed in autoimmune photosensitivity and explains the propensity for inflammatory rather than immunosuppressive responses upon UV exposure in a high type I IFN environment (Fig. 7).

**Figure 7.**
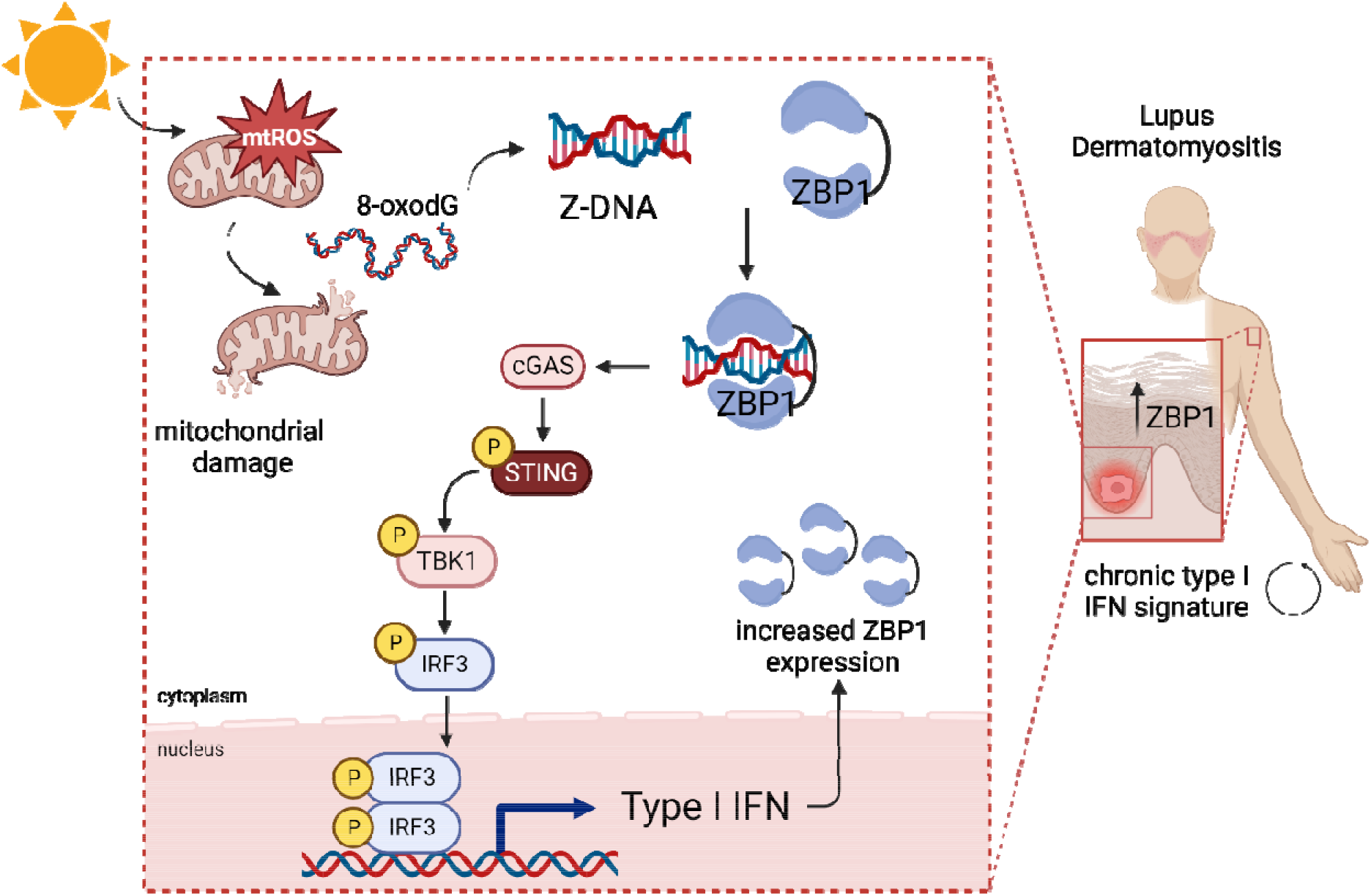
Graphical abstract. UV light promotes mitochondrial ROS formation and mitochondrial damage which results in release of oxidized DNA that can promote Z-DNA conformation. In lupus and dermatomyositis skin, Z-DNA i stabilized by ZBP1 and further activates the cGAS-STING-TBK1-IRF3 pathway to promote type I IFN secretion. This results in more ZBP1 expression, explaining the autocrine loop of type I IFN that i observed in photosensitive autoimmune diseases after UV light.

## Discussion

Here, we describe a novel pathway contributing to photosensitive responses in autoimmune patients. In KCs, chronic IFNα exposure results in upregulation of ZBP1, which stabilizes UVB-induced immunostimulatory Z-DNA derived from oxidized mitochondrial DNA leading to STING-dependent type I and type III IFN production. Knockdown of ZBP1 abrogated IFN responses in KCs and overexpression of ZBP1 resulted in spontaneous IFN activation and Z-DNA accumulation in the cytoplasm. Indeed, our data support a growing list of studies that implicate mitochondrial DNA sensing in a variety of autoimmune diseases including SLE and DM; here, we add skin responses to the list of affected organs and provide the first description for a role of ZBP1 in this process(*26–29, 31, 32*).

The effects of mtDNA oxidation and their contributions to SLE are numerous. Our data show that oxidative DNA lesions after UV exposure promote Z-DNA conformation. In lupus, oxidized mtDNA promotes IFN signaling in neutrophil extracellular traps(*28, 29*). Furthermore, oxidized mtDNA induces a lupus like phenotype in mice, and mtDNA derived from mitochondria-containing red blood cells drives IFN signaling in lupus monocytes(*31, 47*). Whether Z-conformation contributes to these phenotypes will require additional study. Oxidation of genomic DNA confers resistance to degradation by TREX1 and potentiates immunostimulation(*23*). Further, oxidized DNA in the skin of lupus patients colocalizes with the ISG MX1 but evidence that lupus skin shows higher amounts of oxidative DNA lesions is lacking. Our data do not support a role for type I IFN in promotion of ROS in keratinocytes. Rather, our model suggests that oxidized DNA damage by UV results in Z-DNA conformation and that stabilization of the Z-DNA by the ISG ZBP1 is critical to promote more inflammation in diseases such as SLE and DM.

Morphological changes in mitochondria have been associated with mtDNA damage, mitophagy and recently, mitochondrial fragmentation due to apoptotic stress was associated with mtDNA release during senescence(*48*). In our data, we observed a correlation of cytosolic Z-DNA accumulation and mitochondrial fragmentation, indicating mitochondrial dynamics may regulate Z-DNA signaling. It is therefore intriguing to hypothesize that Z-DNA accumulation could also be involved in mtDNA-driven inflammatory states such as cellular senescence.

Our data also prompt the question of whether oxidized mtDNA from other cell types, including immune cells, is likely to transition to Z-DNA and whether Z-DNA sensing may be a central mechanism of the disease-promoting IFN signaling in SLE. Upregulation of ZBP1 is not only observed in lupus skin but also in peripheral blood of SLE patients(*49*). Whether this upregulation is functionally relevant and regulates responses to oxidized mtDNA in immune cells should be further investigated.

Autoantibodies against nucleic acids are a hallmark of SLE development(*50*). Interestingly, SLE patients exhibit autoantibodies against Z-DNA; this was described many years ago and confirmed recently(*51, 52*). However, the substrates for generation of these antibodies have not been studied. Given our data, it is intriguing to propose that Z-DNA derived from UV-irradiated keratinocytes could also serve as an autoantigen and drive the adaptive immune response, especially in the context of UV-mediated systemic disease flares.

Epidermal type I IFN secretion including IFNβ and IFNκ was shown to drive immune responses and enhanced cell death in the skin after UVB exposure(*11*). Other previous studies on mechanisms of photosensitivity in lupus patients were performed in monogenic lupus where cGAS-dependent IFN signaling has been described in TREX1 deficiency(*23, 25*). Nuclear DNA fragments can be released during DNA repair processes and release of nuclear DNA due to rupture of the nuclear envelope can further activate cGAS dependent IFN expression(*53–55*). This has been described in senescence and genetic instability syndromes such as Bloom syndrome(*53, 55–57*). Whether nuclear DNA contributes to STING signaling in autoimmune photosensitivity seems unlikely based on our data and others’. First, the rate of skin cancer in autoimmune photosensitive diseases is not comparable to those with DNA repair deficiencies, indicating no persistence of increased nuclear DNA damage(*58*). Second, type I IFN may increase repair of cyclobutane pyrimidine dimers, the main nuclear DNA lesion upon UV light exposure(*59*). Third, mtDNA is more susceptible to oxidative modifications due to close proximity to ROS production, a lack of protective histones and insufficient DNA repair mechanisms compared to nuclear DNA(*19, 22, 60*). Fourth, we did not observe enhanced Z-DNA formation within the nucleus nor in micronuclei after UV exposure, which can activate cGAS during genetic instability(*61*). Finally, depletion of mitochondrial DNA abrogated UVB-mediated IFN upregulation.

We identified a critical role for ZBP1 in Z-DNA sensing and stabilization after UV exposure. ZBP1 is involved in multiple cellular processes in both antiviral defense via activation of the NF-κB pathway and induction of PANoptosis(*36, 37, 62–65*). Furthermore, it can sustain IFN signaling after release of mitochondrial Z-DNA after Doxorubicin(*34, 35*). So far, previous investigations utilized mostly murine data and investigated cell death pathways: Within the skin, murine ZBP1 was shown to drive necroptosis and autoinflammation only in RIPK1 deficient mice(*39*). In presence of RIPK1, necroptotic signaling of ZBP1 is inhibited and recent evidence revealed that ZBP1 interacts with RIPK1 to sustain IFN signaling in a STAT1 dependent manner(*34, 39*). In our model, cells with overexpression of ZBP1 displayed a spontaneous type I IFN signature without a significant increase of cell death at baseline. We surmise that this may indicate that in an interferon-rich environment, as is seen in autoimmune photosensitive diseases, the primary role of ZBP1 is to bind ZDNA and regulate IFN signaling. It is possible that a second stimulus (such as inhibition of caspase-8 or downregulation of RIPK1) is needed to confer cells towards a cell death phenotype rather than innate immune activation. Previous data of our laboratory found that indeed, IFNα priming promotes apoptotic death rather than other cell death pathways (Loftus *et al.,* under review). Thus, the IFN milieu could act as a “switch” which toggles the function of ZBP1 towards IFN secretion and apoptosis and away from panoptotic cell death. In contrast, diseases with increases in IFNγ show enhanced necroptosis of keratinocytes(*66*). Of note, ZBP1 can also activate the inflammasome via interaction with AIM2, ASC and pyrin to result in IL1β secretion(*67*). Further evidence is needed to understand the role of ZBP1 in inflammasome and cell death pathways in the skin.

Other photosensitive disorders not examined in this paper include porphyrias which are driven by the accumulation of porphyrins due to deficiencies in enzymes involved in hemoglobin metabolism(*68*). How exactly tissue damage and photosensitivity occur in porphyria is currently not understood. Intriguingly, it was shown that ROS accumulate in porphyria and surprisingly, certain porphyrins can stabilize Z-DNA(*69*). This raises the question whether cutaneous Z-DNA might be involved in the entire spectrum of photosensitivity, not just those promoted by autoimmune diseases.

Together, our results uncover a new mechanism of autoimmune photosensitivity driving innate immune responses in the skin of lupus and DM patients that could be important for other photosensitive skin diseases. Both Z-DNA and ZBP1 represent new cutaneous targets for the prevention and potential treatment of lupus and DM skin disease.

## Methods

### Human Subjects

All human subject protocols were reviewed and approved by the University of Michigan IRB-Med. Skin samples from patients with SLE with a history of cutaneous lupus and sex and age-matched healthy controls were obtained with written, informed consent according to the Declaration of Helsinki.

### Cell culture

Immortalized N/TERT keratinocytes (N/TERT-2G) (41), were used with permission from James G. Rheinwald (Brigham and Women’s Hospital, Boston, Massachusetts, USA). N/TERTs were grown in Keratinocyte-SFM medium (ThermoFisher #17005-042) supplemented with 30 μg/ml bovine pituitary extract, 0.2 ng/ml epidermal growth factor, and 0.3 mM calcium chloride. N/TERTs were used from passage 6 to passage 20. Primary human keratinocytes from SLE patients and age and sex matched controls were isolated from non-lesional, non-sun-exposed skin as previously described(*13*) and used at passages 2-6 and grown in Epilife medium (Gibco, #MEPI500CA) with added human keratinocyte growth supplement (10ul/ml medium). Demographics and clinical characteristics of patients and controls used for cell culture are shown in Supplemental Table 1.

STING KO keratinocytes were generated as previously described(*70*). N/TERTs overexpressing *ZBP1* were generated as follows: Full length human Z-DNA binding protein 1 (ZBP1) cDNA, transcript variant 1, was obtained from GenScript (OHu21369). PCR was performed to add the 3XFlag tag to the N terminus of ZBP1 cDNA at the 5’ Not1 site and 3’ Sal1 site to facilitate cloning into expressing vector p3XFlag-CMV-7.1 (Sigma). The resulting plasmid containing the 3XFlag-ZBP1 was further subcloned in the Bst1 and BamHI sites of the vector pLVX-EF1α-AcGFP1-C1 (Takara, cat# 631984, simplified as pLvx-GFP) by PCR to generate the lentivirus overexpressing vector pLVX-GFP-3XFlag-ZBP1 (simplified as pLvx-GFP-ZBP1OE). The constructs were confirmed by sequencing. For Lentiviral infection, the lentiviral empty vector pLvx-GFP and overexpressing pLvx-GFP-ZBP1OE were transiently transfected to 293T cells with packaging plasmids pxPAX2 and pMD2 by the Lipofectamine 2000 to produce the lentivirus as described previously (Bin Xu Plos One 2010). The supernatant containing the lentivirus was used to infect the N/TERTs followed by puromycin selection at 10ug/mL.

Knockdown of ZBP1 in N/TERTs was performed using a lentivirus expressing either shRNA targeting Human ZBP1 or shcontrol. Two different Mission lentivirus-based plasmids of shRNAs (clone numbers TRCN0000123050 and TRCN0000436778) against human ZBP1 and the shcontrol vector TRC2 pLKO.5-puro nonmammalian shRNA (SHC202) were obtained from Sigma-Aldrich (Burlington, MA). 293T cells were cotransfected with the shRNA and packaging plasmids psPAX2 and pMD2 using Lipofectamine 2000 (Invitrogen) in OptiMEM (Gibco) for 6 hrs followed by replacing with keratinocyte medium to produce the lentivirus. Twenty-four hours post transfection, the virus-containing media was collected and centrifuged for 5min at 2000 rpm at 4°C. The resulting supernatant was supplemented with 8 µg/ml Polybrene (Sigma) and used to infect the sub-confluent N/TERTs. The 293T cells were replaced with fresh keratinocyte media and were subsequently used to repeat the N/TERT cell infection two additional times at intervals of 8 to 12 h. N/TERTs infected with either shRNA or shcontrol were selected at 10-12ug/ml puromycin and cells were maintained in 10ug/ml puromycin until the day of experiment.

### UV irradiation

Keratinocytes were grown on either 6 well plates on uncoated glass slides (ThermoScientific, 22×22mm, #3406) for microscopy or 12 well plates for analysis of gene expression. On the day of UVB irradiation, media was changed to prewarmed PBS and cells were irradiated with a dose of 50mJ/cm^2^ using the UV-2 ultraviolet irradiation system (Tyler Research). Emission of UVB radiation (310nm) was allowed by cascade-phosphor ultraviolet generators. Immediately after irradiation, fresh prewarmed media was added until further analysis.

### mitoTEMPO treatment

To reduce mtROS, cells were incubated with mitoTEMPO (Sigma Aldrich, SML0737) (50μM in DMSO) or DMSO 0.5% alone as a control 45min prior to UVB exposure in Keratinocyte SFM. After UVB exposure, fresh mitoTEMPO and medium was added to the wells until further analysis. No differences in mitoSOX staining, confocal microscopy or gene expression were observed with phenol-red containing Keratinocyte SFM vs. phenol-red free media, hence phenol-red containing Keratinocyte SFM was used. Rotenone was used as a positive control for mitoROS (0.5μM).

### Analysis of mitochondrial superoxide (mitoSOX) staining

Keratinocytes were first plated on glass-bottom (no. 1.5) 96-well Mat-Tek dishes and grew for 36h. Cells were then pretreated with mitoTEMPO 45min prior to UVB exposure and then irradiated in PBS (see mitoTEMPO treatment and UV irradiation) and incubated for 30min. Within the last 20 min of the experiment, cells were stained with 2.5μM MitoSOX red dye (Thermo Fisher Scientific, M36008) and Hoechst33342 (1 μg/ml) for 20 min at 37°C, protected from light. Then, cells were washed with PBS and fixed with 4% PFA at RT for 15 min. Lastly, cells were washed with PBS and then immediately imaged using a Nikon Yokogawa X1-CSU spinning disk confocal microscope. Fields of view were selected based on the Hoechst stain.

### RNA isolation, cDNA synthesis and qtPCR

For RNA isolation, the Qiagen RNeasy Plus Mini kit was used according to the manufacturer’s instructions.After isolation of RNA, samples were diluted in 20-40ul RNase free water, depending on target RNA concentrations desired. Total RNA quantification and RNA purity were determined based on the ratio A260nm/A280 nm using Thermo Fisher Scientific NANO drop 2000. A total of 500 to 1000ng of cDNA was synthesized using the iScript cDNA synthesis kit (BioRad, #1708891). 10ng cDNA was used for quantitative PCR in technical triplicates in a 384 well plate using SYBR Green Supermix (Applied Biosystems). PCR was run in Applied Biosystems QuantStudio™ 12K Flex Real-Time PCR at the Advanced Genomics Core at the University of Michigan. Primer sequences are summarized in Supplemental Table 4. Gene expression level was determined by relating to the housekeeper gene beta-Actin or RPLP0 using the ΔΔct method setting the mock condition to one.

### Immunofluorescence staining cell culture

Keratinocytes were plated on coverslips in a 6 well plate. Mitochondrial ROS were stained 30min after UVB exposure. 8oxodG, Z-DNA and mitochondrial dynamics were stained 3h after UVB exposure. At each experimental endpoint, cells were fixed with freshly prepared 4% paraformalaldehyde (PFA) at RT for 15min. The IFA was always performed on the same day as the experiment. After fixation, cells were washed with PBS + 0.1% TritonX100 (wash buffer) and then blocked with 5% BSA and 10% normal goat serum in wash buffer (block buffer) for 30min at RT. Primary antibody cocktails were prepared in block buffer and incubated on each coverslip for 1h at RT. Then, wells were washed three times using wash buffer and incubated with secondary antibodies and counterstains diluted in block buffer for 30min at RT. After this, wells were washed three times using wash buffer and coverslips were mounted on glass slides using ProLong Glass Antifade Mountant (Invitrogen, P36980). Large (3×3) images were taken the next day either with 60X or 100X magnification, depending on the staining, on a Nikon Yokogawa spinning disk microscope. Using large image acquisition, over 250 - 500 cells per condition could be stained. Specific antibody details are available in Supplemental Table 3. Operator bias was reduced during image acquisition through selection of fields of view and focal planes based on counterstains that were unrelated to the experimental question (DAPI or CellTracker).

### Intensity Line Measurement

Representative images for intensity measurement were taken on a Nikon Yokogawa spinning disk microscope and pixel intensity was assessed with Fiji (ImageJ) to measure indicated staining (TOMM20, Z-DNA and 8oxodG) intensity across the dotted lines at baseline and after UVB exposure.

### Automatic image analysis

Confocal microscopy images were quantified by automated image analysis using CellProfiler, an open-source software. All CellProfiler pipelines associated with this work are available in supplemental files (Supplemental file 1). Analysis of images was performed on raw images unless otherwise specified. Briefly, single-cells were identified by nuclear objects based on global thresholding of nuclear staining (DAPI, 4’,6-diamidino-2-phenylindole or Hoechst) using the “identify primary objects” module, followed by propagation of the nuclear objects to the cellular periphery based on a whole-cell stain (CellTracker) using the identify secondary objects module. Subsequently, a variety of cellular parameters were measured and related to parent cells using the relate objects module.

For the quantification of mitochondrial superoxide (Supplemental file 1, Figure 1 C), the intensity of the mitochondrial superoxide indicator MitoSOX was analyzed within cells using Hoechst as the nuclear counterstain and defining cells by propagation of the nuclear objects to the cellular periphery based on mitoSOX staining.

Mitochondria were determined using the TOMM20 immunostaining with two-class Otsu adaptive thresholding within the “identify primary objects” module, and the mitochondrial compartment was determined by using the “merge objects” to connect adjacent mitochondrial objects (neighbor distance 0 pixels). To measure Z-DNA puncta outside of the mitochondrial compartment (Supplemental file 1, Figure 1L), puncta in the Z-DNA immunostaining were identified using the primary objects module. The cytosolic mitochondria-free content was created using the “identify tertiary objects” module by defining the non-nuclear compartment as the CellTracker-positive and DAPI-negative space and subsequently the cytosol by subtracting the mitochondrial staining (by TOMM20) from the non-nuclear compartment to get the mitochondria-free cytosolic compartment. Tertiary objects were then related to cells by using the “relate objects” module to determine cytosolic Z-DNA puncta per cell.

For analysis of 8-Oxo-2’desoxyguanosine (8oxodG, Supplemental file 1, Fig. 2, A and D), the intensity of the 8oxodG staining was analyzed within cell objects (defined as ‘total’ in the manuscript) as well as within the mitochondrial compartment which was defined by segmentation of mitochondrial staining (anti-TOMM20). For the correlation of 8oxodG with total and cytosolic Z-DNA (Figure 2D), intensity values of 8oxodG per cell and Z-DNA puncta per cell across all images and different conditions were matched and the average intensity of 8oxodG for the counts of Z-DNA puncta was calculated. The corresponding intensity to Z-DNA puncta counts were then plotted and correlated using Pearson correlation analysis. For correlation of mitochondrial fragments with total and cytosolic Z-DNA, mitochondrial fragments per cell were calculated, plotted against Z-DNA puncta counts per cell and then correlated using Pearson correlation analysis.

In Figure 4F, intensity values for epidermal ZBP1 were determined using manual identification of the epidermis based on nuclear staining of slides with DAPI. Average intensity per image with similar epidermal areas including >300 cells were included in the analysis.

In Figure 6G, cGAS intensity per cell was quantified within nuclear objects, and the total cytoplasm (nuclear subtracted cellular area, based on GFP). The the nuclear:cytoplasmic ratio was calculated by dividing nuclear and cytoplasmic intensity for each cell.

Removal of outliers resulting from automated image analysis was performed using strict ROUT outlier identification (*Q* = 0.1%) in GraphPad Prism from cell-level data pooled across multiple experiments. Representative confocal images shown in this manuscript were prepared using the ImageJ background subtraction tool with a rolling ball radius of 30 pixels. Operator bias during image acquisition was reduced by selecting fields of view and cells based on counterstains that were unrelated to the staining of primary interest.

### Depletion of mtDNA using ddC

To reduce total mtDNA content, we treated N/TERTs with 2′3′-dideoxycytidine (ddC) according to a previously published protocol(*71*) with the following modifications: N/TERTs were seeded in a 6 well plate at 15-20% confluency and 12h after seeding medium was changed containing ddC (50μM and 150μM). Medium was changed every 24h until confluency reached 70-80% after 2 days. Then, cells were irradiated with UVB exposure (see above). Z-DNA staining was assessed 3h after UVB exposure and gene expression was assessed 6h after irradiation.

### Cellular ROS measurement

For detection of cellular ROS, we used the indicator CM-H_2_DCFDA (Invitrogen) and cells were incubated with CM-H_2_DCFDA 10uM for 30 minutes prior to UVB exposure. Phenol red-free keratinocyte media (Gibco) was added, and fluorescence was measured five minutes post UVB exposure using a microplate reader (Ex/Em 492/527 nm). The average of 3-5 replicates per condition were averaged in each experiment (n=4-6), background fluorescence was subtracted, and data was expressed as fold change relative to untreated.

### Measurement of B-DNA and Z-DNA conformation using ratio of A260/295

Conformation of Z-DNA and B-DNA was assessed using the absorbance ratio of 260 to 295nm as previously described(*43, 45*). We used poly(dGdC) (50ng/µl) for conformation analysis. Poly(dGdC) was diluted in water with increasing salt concentrations (titration of 1.8M, 2M, 2.2M, 2.4M, 2.6M, 2.8M, 3M, 3.2M, 3.4M, 3.6M and 4M NaCl) to induce Z-DNA conformation. Reaction was performed for 2h at 37°C in nuclease-free microcentrifuge tubes (BioRad). A260/295 was measured with a nanodrop (company) using water with different salt concentrations as a blank. For induction of Z-DNA by H2O2, poly(dGdC) was diluted in water containing 1mM H2O2 and increasing salt concentrations as mentioned above. Samples were incubated for 2h at 37°C and were measured with a nanodrop using 1mM H202 with different salt concentrations (absent DNA) as blanks. Each incubation was performed in triplicates. The values were then plotted as the ratio of A260/295.

### Skin genome-wide expression datasets

Three previously published microarray datasets were used for analysis. These included samples from: 1. healthy controls (n=13) and lesional skin of patients with lupus (n=90, of these 47 DLE and 43 SCLE)(*72*); 2. healthy controls (n=5) and dermatomyositis (n=41 biopsies from 36 patients)(*16*); and 3. healthy pediatric controls (n=8) and childhood onset systemic lupus (n=7 lesional skin biopsies from 5 patients) and juvenile dermatomyositis (n=9 lesional skin biopsies from 9 patients)(*73*). Microarray datasets are available from CLE through GEO GSE81071, from adult DM through GSE142807, from childhood SLE and juvenile DM through GEO GSE148810. As we previously published, Pearson correlation analysis was performed between gene expression and a previously described 6 IFN-stimulated gene score(*73, 74*) using GraphPad Prism version 9.1.0.

### Single cell Sequencing

For Fig. 3, F, expression of *ZBP1* was examined from our single cell RNA sequencing dataset in nonlesional (n=14), lesional (n=14) and healthy control (n=14) skin(*12*). Specifically, ZBP1 expression was plotted across the major cell types defined in the single cell RNA sequencing dataset, and these cell types were further divided by the disease states of the cells.

### Tissue immunofluorescence

To assess tissue protein expression, formalin-fixed, paraffin-embedded tissue slides were obtained from patients with cutaneous lupus (chronic discoid lupus, subacute cutaneous lupus and nonlesional skin from systemic lupus patients), dermatomyositis (lesional and nonlesional skin) and healthy controls. Slides were heated for 1h at 60°C, rehydrated, and antigen retrieved with tris-EDTA (pH6). Slides were blocked with blocking buffer (PBS + 10% normal goat serum) and then incubated with primary antibodies (diluted in blocking buffer) against ZBP1 (Supplemental Table 3) overnight at 4°C. Slides were incubated with secondary antibodies (Supplemental Table 3) and counterstained with 4′,6-diamidino-2-phenylindole (DAPI). Slides were mounted with Antifade glass mounting medium (ThermoFisher) and dried overnight at room temperature in the dark. Images were acquired using a Zeiss Axioskop 2 microscope and analyzed using CellProfiler (see above). Images presented are representative of at least five biologic replicates. Patient characteristics for samples from SCLE/DLE and DM patients are shown in Supplemental Table 2. Controls were anonymized biopsies from healthy control without history of skin disease.

### DNA transfection

Primary control keratinocytes or N/TERTs were seeded in triplicates in a 12 well plate and grown for 48h in appropriate media to 60-70% confluence (see above). DNA transfection was performed using Lipofectamine2000 and diluted dsDNA (50ng/μl, B-DNA equivalent) as well as polydGdC (50ng/μl, Z-DNA equivalent). After warming all reagents and media up to RT, Lipofectamine2000 (11668–029) and DNA were diluted in OptiMEM (Gibco, #31985-070) at a ratio of 3μl/μg DNA. Cells were covered with 400μl of OptiMEM containing either 50ng DNA (125ng DNA/ml media) or 100ng DNA (250ng DNA/ml media) and incubated at 37C until further analysis. 4h after transfection, cells were washed with ice-cold PBS and then lysed for phospho-Western Blot with Protease inhibitor (cOmplete Mini, EDTA-free, Sigma, #11836170001) and Phosphatase Inhibitor Cocktail (Pierce, PI78420). 6h and 24h after transfection, cells were analyzed for gene expression using the Qiagen RNeasy Plus Mini Kit and qtPCR (see above).

### Western Blot

Cells were washed with ice-cold PBS and then lysed. Prior to sodium dodecyl sulfate-polyacrylamide gel electrophoresis (SDS-PAGE), sample protein content was normalized by dilution following a Bradford assay. Samples were diluted in Laemmli sample loading buffer supplemented with β-mercaptoethanol (Bio-Rad), heated for 5 min at 95°C, and then separated on 4 to 20% gradient polyacrylamide tris-glycine gels (Bio-Rad). After SDS-PAGE, gels were transferred to a 0.45-μm nitrocellulose membrane by a semi-dry transfer system (Cytiva) and membranes were blocked with PBS + 5% BSA + 0.1%Tween20 (blocking buffer) for 30min at room temperature on a shaker. After blocking, primary antibodies were diluted in blocking buffer and added to incubated overnight at 4°C on a shaker. Secondary antibodies diluted in blocking buffer were added and incubated for 30min at room temperature in the dark. Image acquisition was accomplished using LI-COR IR dye secondary antibodies and an Odyssey IR Imager. Quantification of Western blots was performed using ImageJ densitometric gel analysis for 1D gels. Antibodies and dilutions for Western Blot are listed in Supplemental Table 3.

## Acknowledgements

We thank the patients for participation in this study. We thank Chrissy Goudsmit and Sri Yalavarthi for helpful technical assistance. We acknowledge the assistance of the Microscopy core and the Advanced Genomics core at the University of Michigan.

## Author contributions

Conceptualization: BK, MK; Methodology: BK, MR, BX, MGK, CB, SH, SE, JEG, KEM, AV, CD, MOR, JMK; Investigation: BK, MR, BX, MGK, YG, CB, AV, SE, SH, CD, GH, FM, JT, JEG, MOR, JMK; funding acquisition: BK, MR, MOR, MK; Visualization: BK, MR, BX, MGK, CB, SH; project administration: MK; Supervision: JEG, MOR, JMK; writing-original draft: BK, KM; writing – review and editing: BK, MR, JT, JEG, MOR, JMK

## Data and materials availability

Skin wide genome expression datasets are already available for CLE through GEO GSE81071, from adult DM through GSE142807, from childhood SLE and juvenile DM through GEO GSE14 8810. The scRNA-seq data are available in GEO under accession number GSE186476. Pipelines for automated image analysis in Cell Profiler are within the Supplemental files of this manuscript. All other data supporting the conclusions of the manuscript including patient data are available in the main text or the supplemental figures and tables.

## Supplemental Data

**Supplemental Figure 1.**
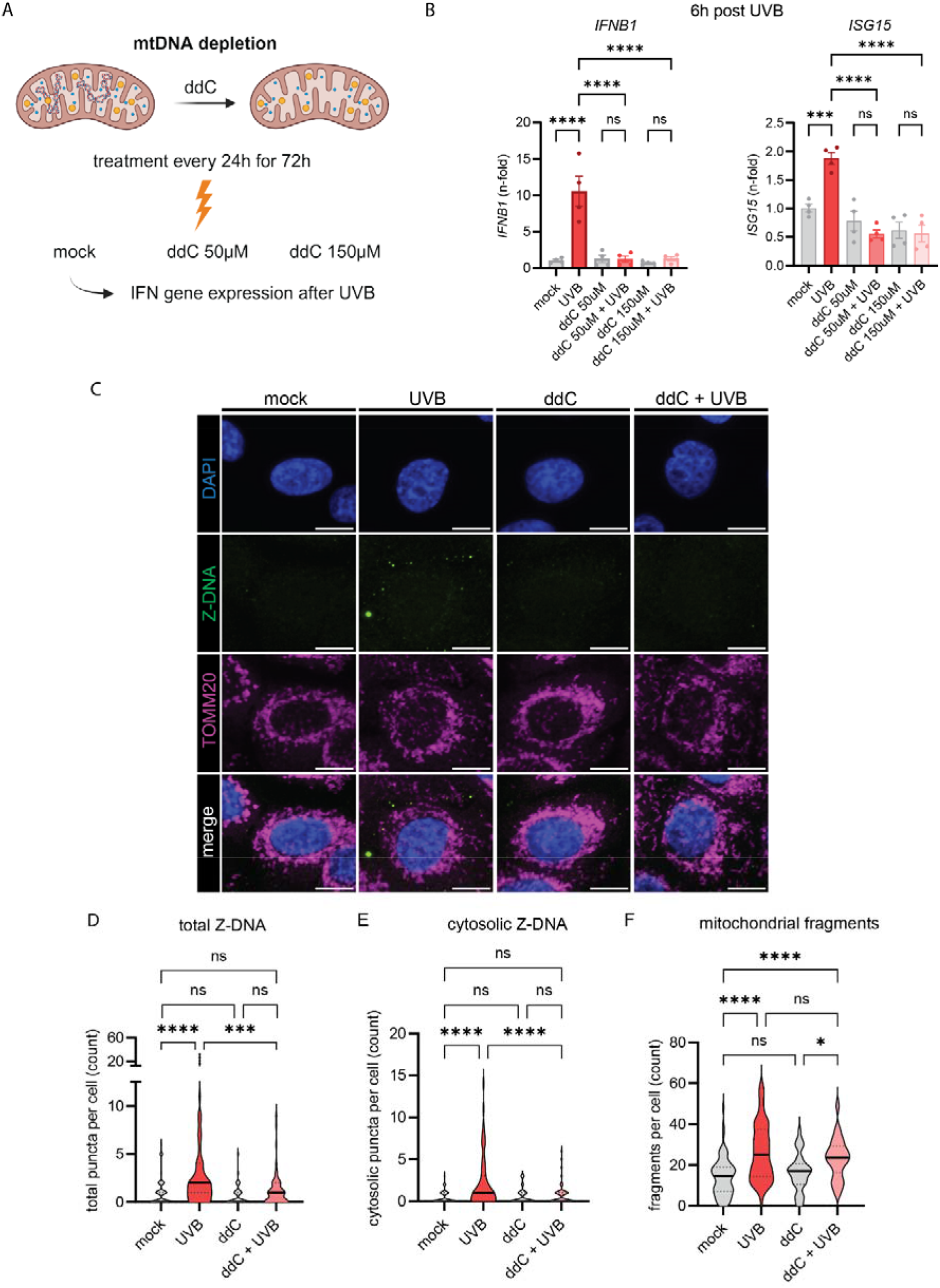
UVB-driven IFN responses are mtDNA dependent and UV-induced Z-DNA derives from mtDNA. **A.** Experimental approach for mtDNA depletion in N/TERTs using nucleoside 2’,3’ dideoxycytidine (ddC). Treatment with ddC was performed for 72h. After irradiation, medium was changed to ddC-free medium until gene expression analysis. **B.** Quantitative gene expression 6h after UVB exposure. n=2 for each experiment. **C.** Representative confocal images of N/TERTs treated with +/-ddC +/-UVB 3h after UVB exposure stained for Z-DNA, TOMM20 and DAPI. Scale bar 10μm. **D.** Quantification of Z-DNA puncta and mitochondrial fragments using CellProfiler open-source software from conditions in (C.), n=3. Comparisons were done via ordinary one-way ANOVA followed by Sidak’s multiple comparison test. Mean and SEM. *P<0.05, **P<0.01, ***P<0,001, ****P<0.0001.

**Supplemental Figure 2.**
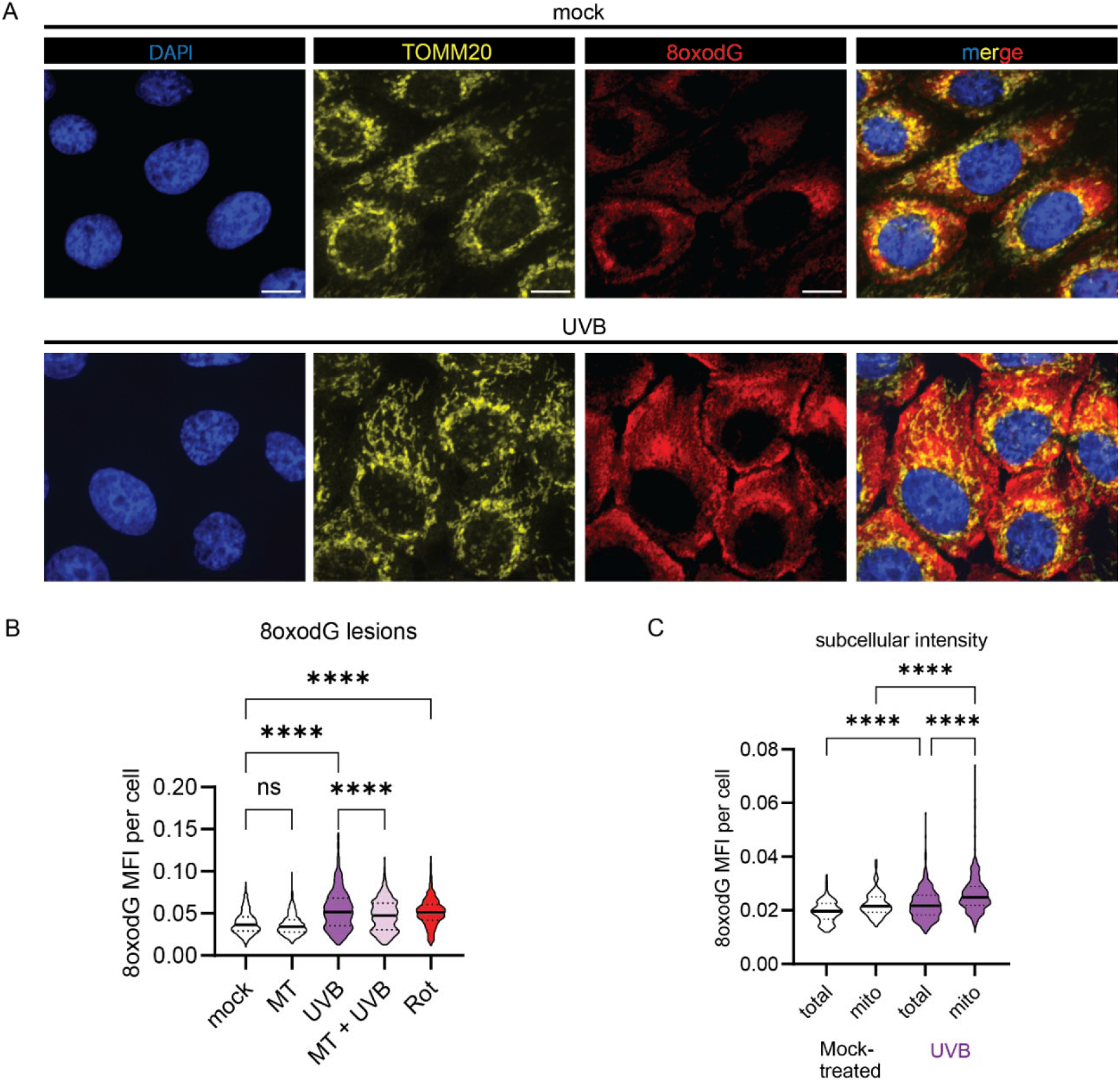
UVB induces oxidative DNA damage in the cytosol and mitochondrial compartment. **A.** Representative confocal microscopy images of N/TERTs 3h after UVB exposure stained for TOMM20, 8oxodG lesions and DAPI. Scale bar 20μm. **B.** Quantification of 8oxodG intensity per cell using open-source software, CellProfiler, in N/TERTs treated +/-mitoTEMPO (50μM), +/-UVB or Rotenone (0.5μM) as a positive control, n=3. **C.** Quantification of subcellular intensity of 8oxodG intensity per cell (total) or mitochondrial (mito) assessed by TOMM20^+^ merged area. Comparisons were done via ordinary one-way ANOVA followed by Sidak’s multiple comparison test. Mean and SEM. *P<0.05, **P<0.01, ***P<0,001, ****P<0.0001.

**Supplemental Figure 3.**
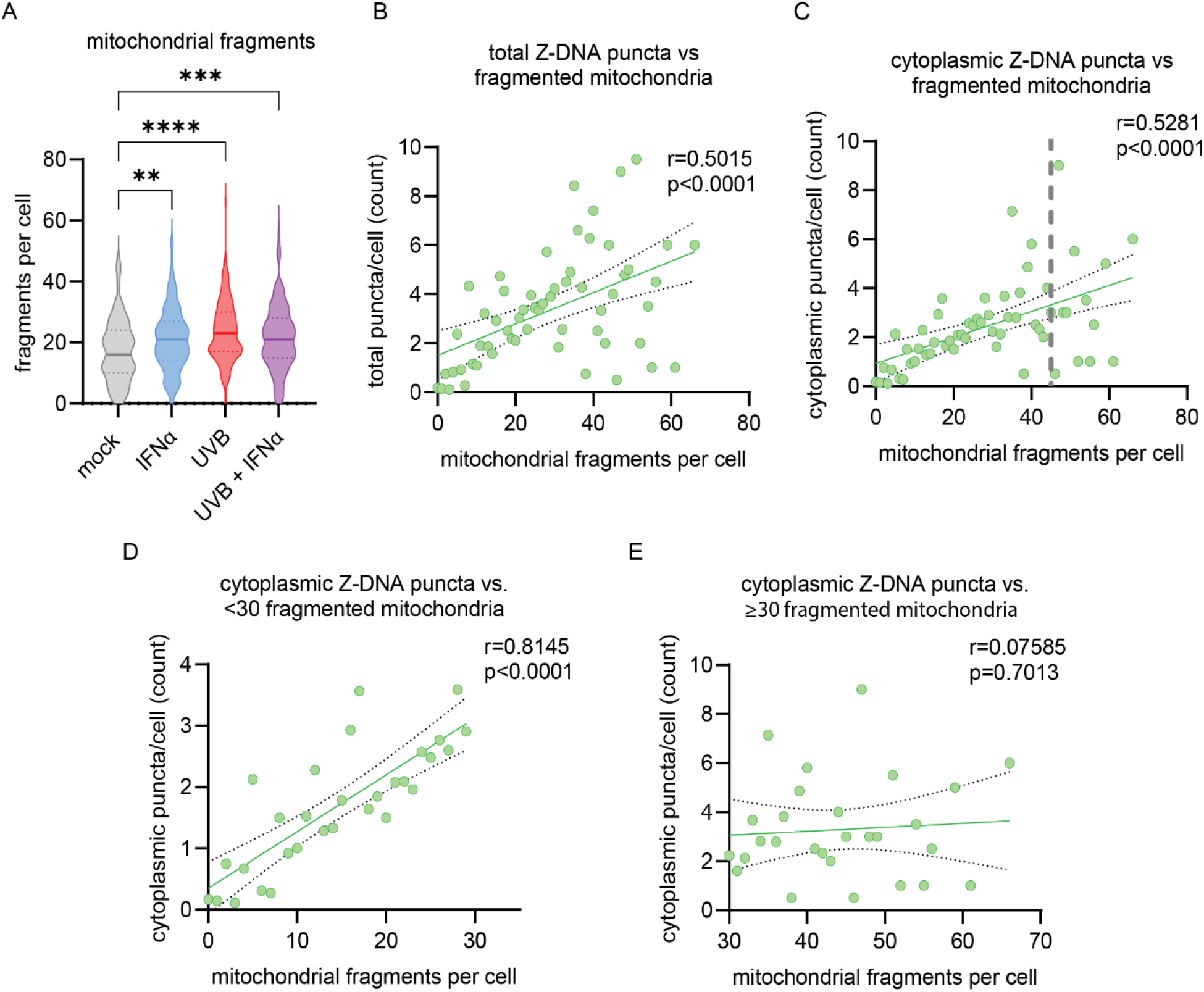
Cytosolic Z-DNA accumulation is associated with mitochondrial fragmentation. **A.** Violin plots represent quantification of mitochondrial fragments (defined as TOMM20^+^ objects smaller than 1μm) in N/TERTs after 16h of IFNα treatment followed by UVB (50mJ/cm^2^) exposure. Comparisons were done via ordinary one-way ANOVA followed by Sidak’s multiple comparison test. **P<0.01, ***P<0,001, ****P<0.0001. B and C. Correlation of total or cytoplasmic Z-DNA puncta and fragmented mitochondria with simple linear regression. D and E. Correlations of data in C divided by # of mitochondrial fragments per cell. Pearson correlation coefficient (r) and p-values for indicated correlations are shown in the upper right.

**Supplemental Figure 4.**
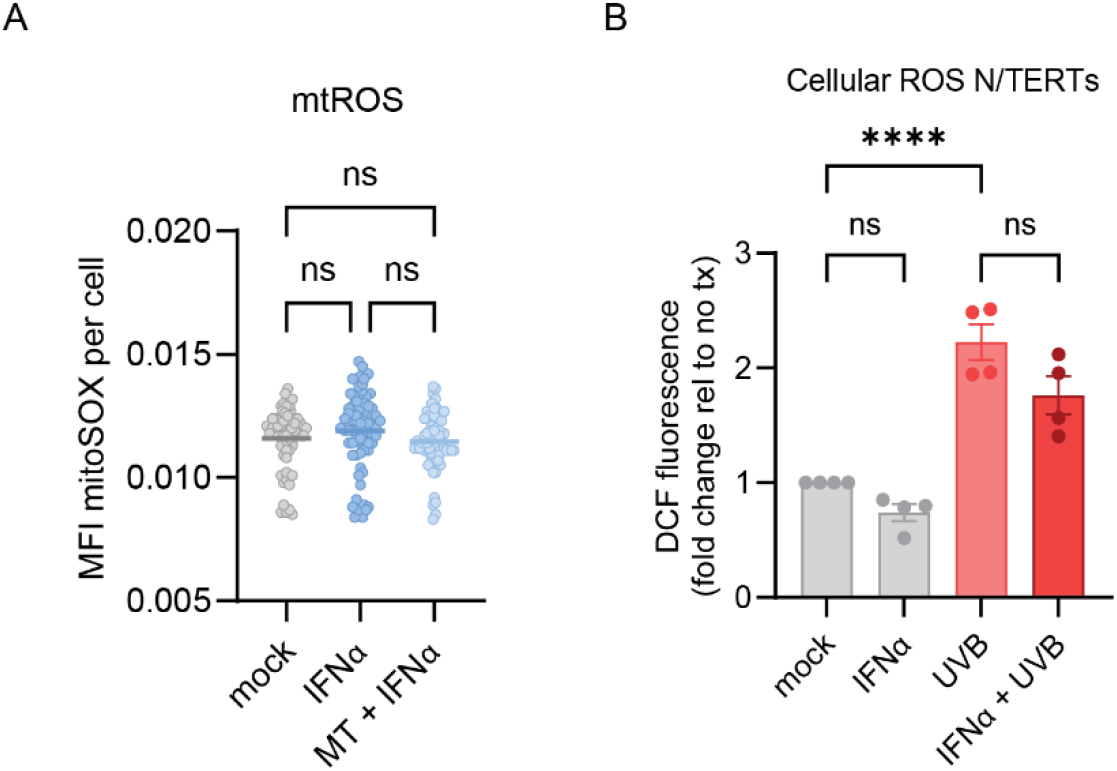
IFNα does not increase mitochondrial or total cellular ROS in N/TERTs. **A.** Violin plots represent quantification of mitoSOX staining intensity per cell in N/TERTs stimulated with IFNα (1000U/ml) for 16h compared to mock. B. Fold change of fluorescence of Dichlorodihydrofluorescein (DFC) after treatment with IFNα for 16h +/-UVB exposure in N/TERTs 5min after UVB exposure, n=4. Comparisons were done via ordinary one-way ANOVA followed by Sidak’s multiple comparison test. Mean and SEM. ****P<0.0001.

**Supplemental Figure 5.**
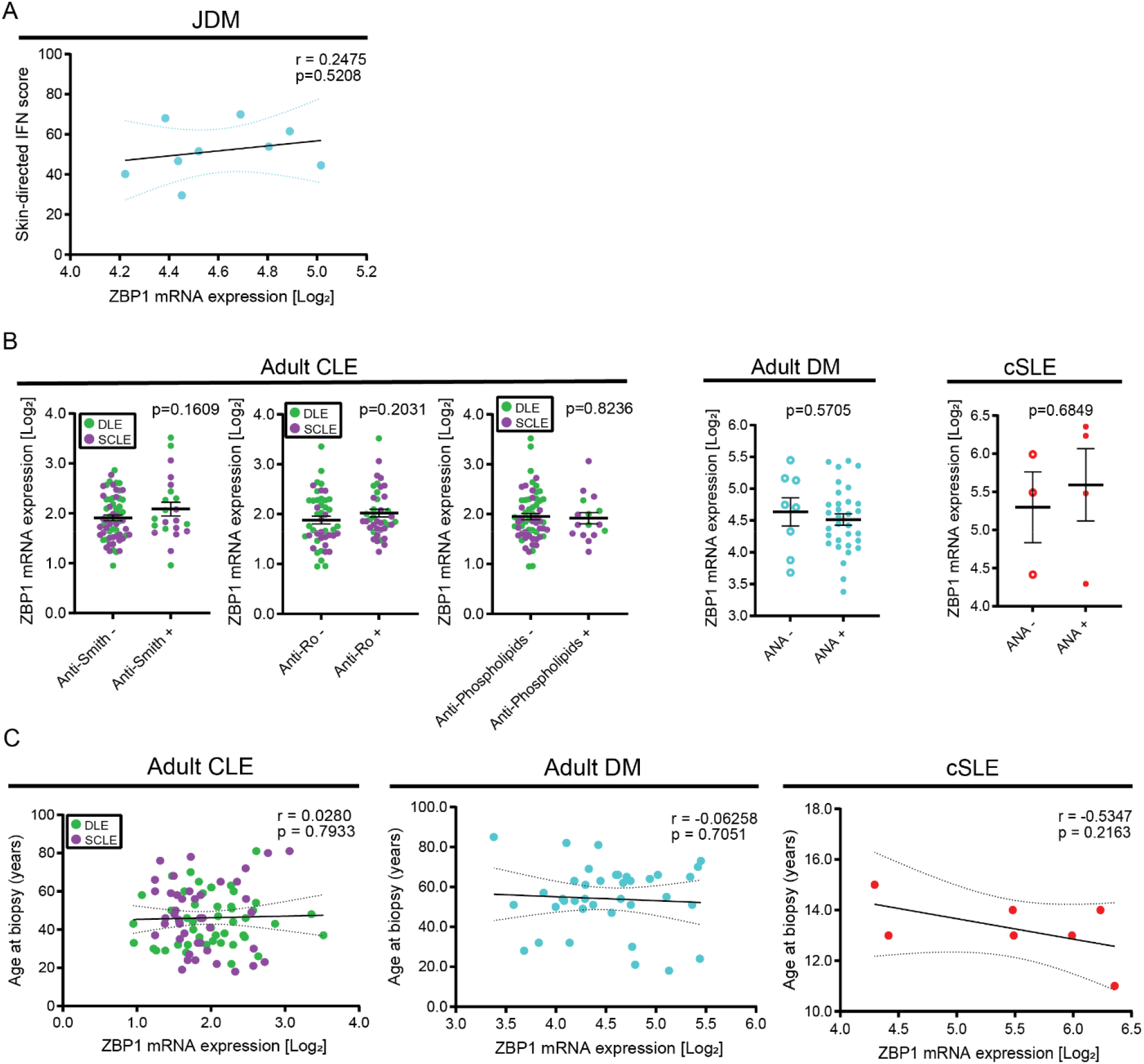
ZBP1 expression does not correlate with systemic autoantibodies or patient age. **A.** Correlation of cutaneous *ZBP1* expression in juvenile dermatomyositis (n=9) with skin-directed IFN score showing no significant correlation. **B.** Comparison of cutaneous *ZBP1* expression with autoantibodies in adult CLE, DM and childhood onset SLE (cSLE) showing independence of *ZBP1* expression with autoantibody status. **C.** Correlation of cutaneous *ZBP1* expression with age in adult CLE, adult DM and childhood SLE (cSLE) showing no significant correlation with age.

**Supplemental Figure 6.**
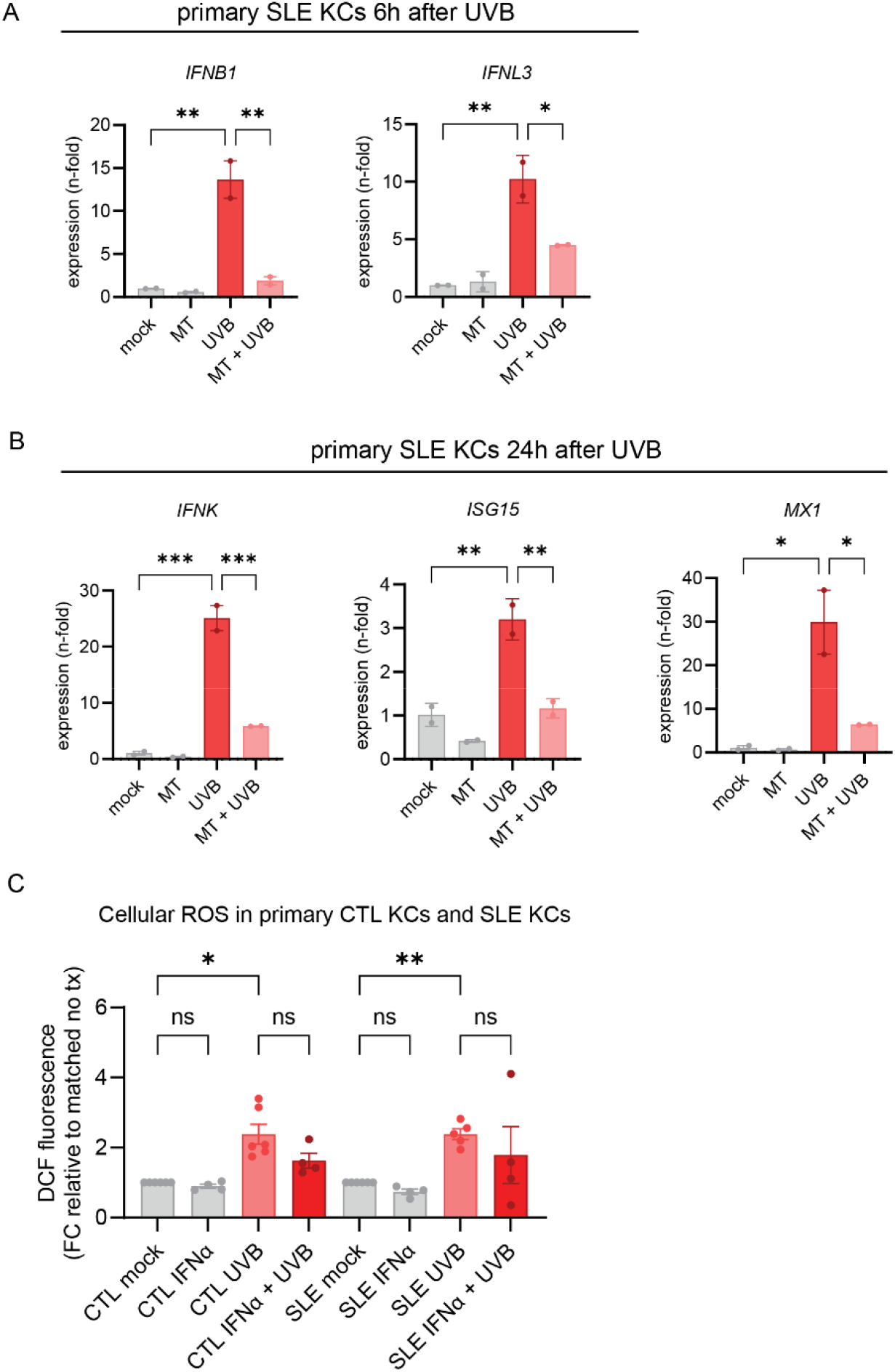
mitoTEMPO rescues UVB-induced IFN expression in lupus KCs. **A.** Nonlesional SLE KCs (n=2) were treated +/-mitoTEMPO (50μM) and irradiated with UVB. Gene expression was analyzed 6h after UVB exposure. **B.** Gene expression analysis 24h after UVB exposure was normalized to β-Actin. n=2. Mean and SEM. **C.** Measurement of cellular ROS in primary HC KCs (n=4) and SLE KCs (n=4) at baseline and after IFNα treatment +/-UVB exposure. Comparisons were done via ordinary one-way ANOVA followed by Sidak’s multiple comparison test. Mean and SEM. *P<0.05, **P<0.01, ***P<0,001, ****P<0.0001.

**Supplemental Figure 7.**
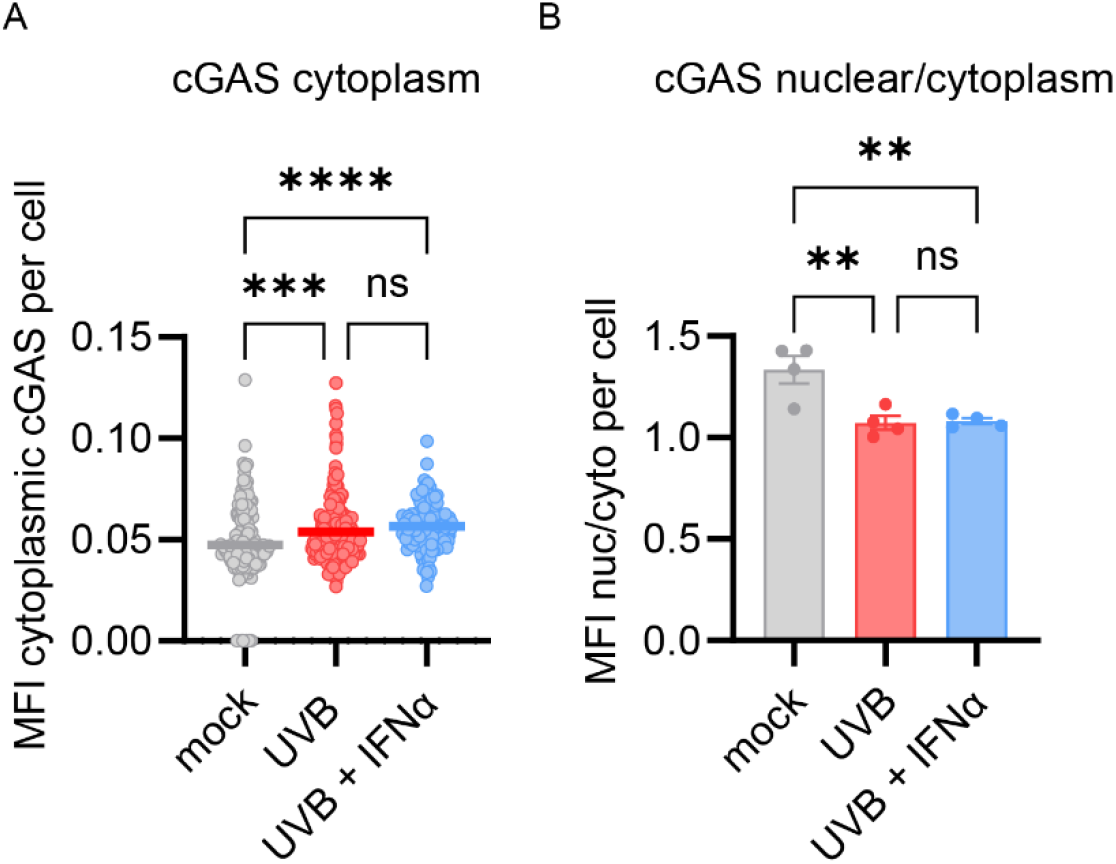
UVB leads to cytosolic shift of cGAS in N/TERTs. **A**. Quantification of cytosolic mean fluorescence intensity (MFI) of cytoplasmic cGAS defined by the DAPI-negative area in N/TERTs using open-source software CellProfiler. **B.** Ratio of nuclear and cytoplasmic MFI per cell and shown is the mean ratio per cell of each experiment (n=4). Comparisons were done via ordinary one-way ANOVA followed by Sidak’s multiple comparison test. Mean and SEM. **P<0.01, ***P<0,001, ****P<0.0001.

**Supplemental Figure 8.**
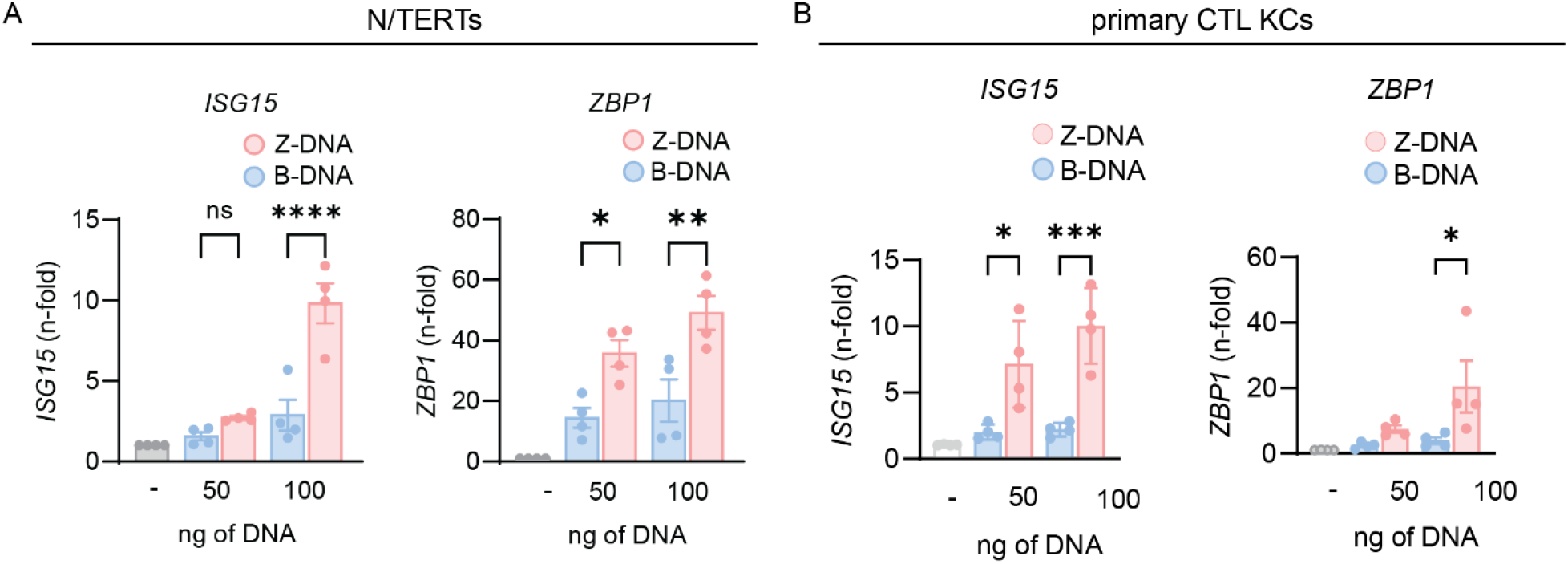
ISG15 and ZBP1 are significantly increased after Z-DNA transfection vs. B-DNA in N/TERTs and primary KCs. **A**. Gene expression at 24h of indicated genes from N/TERTs (n=4) and primary HC KCs (n=4) treated transfected with Z-DNA or B-DNA. Comparisons were done via ordinary one-way ANOVA followed by Sidak’s multiple comparison test. Mean and SEM. **P<0.01, ***P<0,001, ****P<0.0001.

**Supplemental Figure 9.**
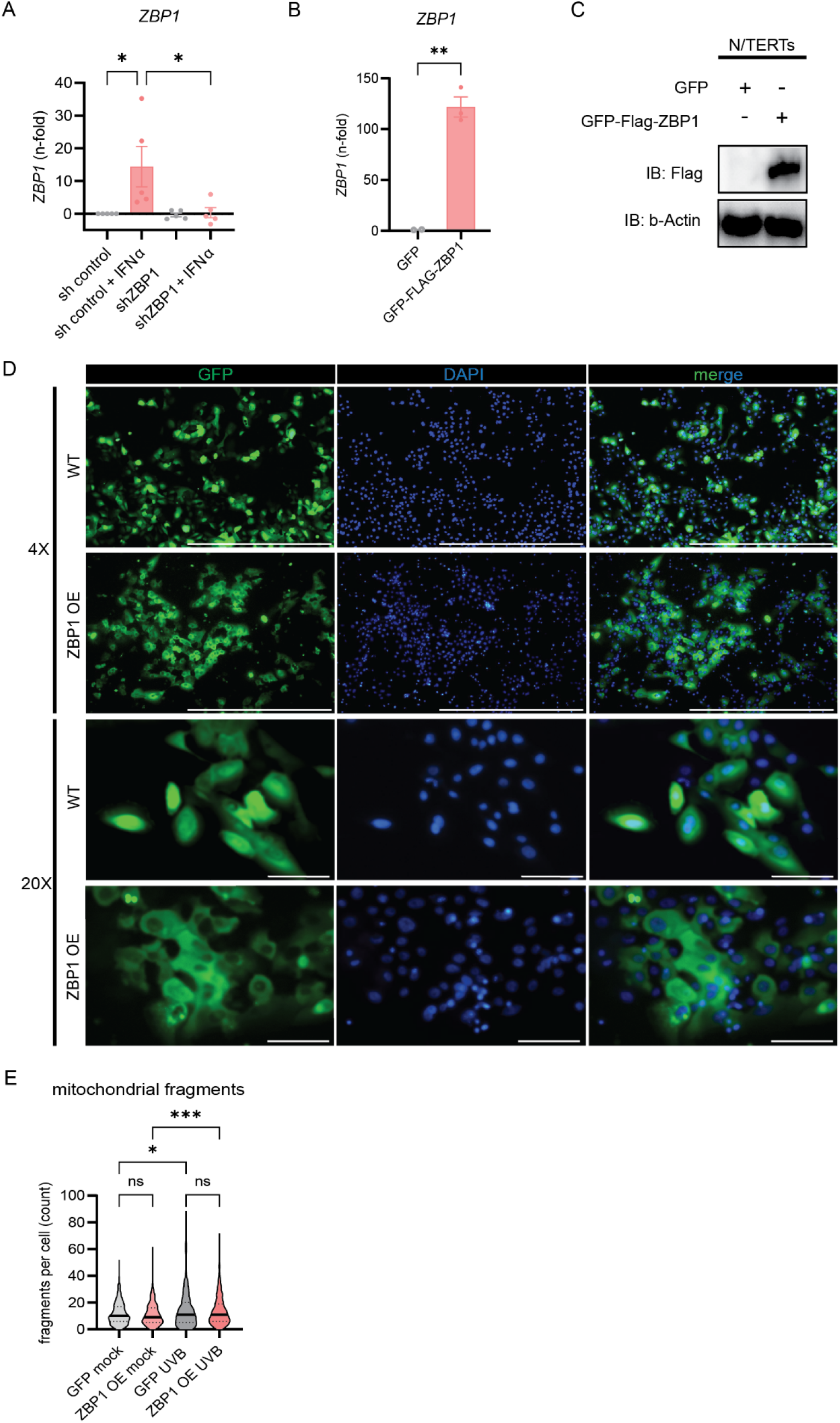
Overexpression of ZBP1 results in cytosolic expression. A. Confirmation of shRNA knockdown by qPCR compared to shcontrol after IFNα stimulation (1000U/ml) for 16h, n=5. B. Quantitative gene expression of ZBP1 overexpressors compared to GFP alone, n=3. C. Immunoblot against FLAG confirming FLAG-tag of ZBP1 overexpressor cells. D. Representative immunofluorescence images show efficient transfection of both GFP (first line) alone and GFP-ZBP1 (second line) in 4X magnification, scale bar=1000μm. Detailed images reveal pancellular tag of GFP (third line) and cytosolic overexpression of ZBP1 (fourth line). 20X, scale bar=1000μm. 4X, scale bar=100μm. E. Quantification of mitochondrial fragments (TOMM20^+^ objects <1µm^2^ with circularity >0.6) in GFP-tag N/TERTs and ZBP1 OE N/TERTs at baseline and after UVB exposure using CellProfiler software. Comparisons were done via ordinary one-way ANOVA followed by Sidak’s multiple comparison test or t-test. *P<0.05, **P<0.01, ***P<0,001, ****P<0.0001.

**Supplemental Table 1.** Demographics and characteristics of patients and controls for primary keratinocyte cell culture

|  | **HC (N=8)** | **SLE (N=8)** |
| --- | --- | --- |
| Median age in years (IQR) | 44 (31,52) | 44 (41,52) |
| Female sex - n (%) | 4 (50%) | 6 (75%) |
| Cutaneous lupus – n (%) | - | 5 (62%) |
| Median SLEDAI-2k (IQR) | - | 2 (0,4) |
| **Cutaneous lupus subtype – n (%)** |  |  |
| ACLE | - | 1 (12%) |
| SCLE | - | 1 (12%) |
| DLE | - | 3 (38%) |
| CLASI activity (IQR) |  | 2 (0,3) |
| **SLE treatment – n (%)** |  |  |
| Hydroxychloroquine | - | 5 (62%) |
| Glucocorticoid | - | 3 (38%) |
| Immunosuppressant | - | 7 (88%) |
| **Autoantibodies – n positive (%)** | - |  |
| ANA | - | 8 (100%) |
| Anti-Ro/SSA | - | 5 (62%) |
| Anti-dsDNA | - | 4 (50%) |
| Anti-Sm/RNP | - | 4 (50%) |
| **Site of non-lesional biopsy - n (%)** |  |  |
| Buttock/hip | 8 (100%) | 7 (88%) |
| Arm | 0 | 1 (12%) |
| HC: healthy controls; SLE: systemic lupus erythematosus; IQR: interquartile range; n: number; SLEDAI: Systemic Lupus Erythematosus Disease Activity; ACLE: acute cutaneous lupus; DLE: discoid lupus erythematosus; SCLE: subacute cutaneous lupus erythematosus; CLASI: Cutaneous Lupus Erythematosus Disease Area and Severity Index; ANA: antinuclear antibody |  |  |

**Supplemental Table 2.** Demographics and characteristics of lupus and dermatomyositis patients from which skin biopsies were used for tissue immunofluorescence

|  | CLE/SLE (N=13) |  | DM (N=6) |  |
| --- | --- | --- | --- | --- |
| Median age in years (IQR) | 46 (41,51) |  | 54 (35,61) |  |
| Female sex - n (%) | 11 (85%) |  | 5 (83%) |  |
|  | Clinical manifestations CLE/SLE |  | Clinical manifestations DM |  |
|  | Cutaneous lupus only – n (%) | 4 (31%) | Skin involvement | 6 (100%) |
|  | DLE | 8 (62%) | Muscle involvement | 4 (67%) |
|  | SCLE | 5 (38%) |  |  |
|  | Median CLASI activity (IQR) | 4 (2,8) |  |  |
|  | Median SLEDAI-2k (IQR) | 4 (2,8) |  |  |
| Autoantibodies – n positive (%) |  |  |  |  |
| ANA | 12 (92%) |  | 5 (83%) |  |
|  | Anti-Ro/SSA | 6 (46%) | Anti-Mi-2 | 1 (17%) |
|  | Anti-dsDNA | 4 (31%) | Anti-TIF-1γ | 1 (17%) |
|  | Anti-Sm/RNP | 3 (23%) | Anti-PL7 | 1 (17%) |
| Treatment – n (%) |  |  |  |  |
| Hydroxychloroquine | 12 (92%) |  | 2 (33%) |  |
| Glucocorticoid | 6 (46%) |  | 1 (17%) |  |
| Immunosuppressant | 5 (38%) |  | 3 (50%) |  |
| SLE: systemic lupus erythematosus; CLE: cutaneous lupus erythematosus; DM: dermatomyositis; IQR: interquartile range; n: number; DLE: discoid lupus erythematosus; SCLE: subacute cutaneous lupus erythematosus; CLASI: Cutaneous Lupus Erythematosus Disease Area and Severity Index; SLEDAI: Systemic Lupus Erythematosus Disease Activity; ANA: antinuclear antibody |  |  |  |  |

